# Phenylhydrazone-based Endoplasmic Reticulum Proteostasis Regulator Compounds with Enhanced Biological Activity

**DOI:** 10.1101/2025.04.04.646800

**Authors:** Gabriel M. Kline, Lisa Boinon, Adrian Guerrero, Sergei Kutseikin, Gabrielle Cruz, Marnie P. Williams, Ryan J. Paxman, William E. Balch, Jeffery W. Kelly, Tingwei Mu, R. Luke Wiseman

**Affiliations:** Department of Chemistry, The Scripps Research Institute, La Jolla, CA 92037; The Department of Physiology and Biophysics, Case Western Reserve University, Cleveland, OH 44106; Department of Molecular and Cellular Biology, The Scripps Research Institute, La Jolla, CA 92037

## Abstract

Pharmacological enhancement of endoplasmic reticulum (ER) proteostasis is an attractive strategy to mitigate pathology linked to etiologically-diverse protein misfolding diseases. However, despite this promise, few compounds have been identified that enhance ER proteostasis through defined mechanisms of action. We previously identified the phenylhydrazone-based compound AA263 as a compound that promotes adaptive ER proteostasis remodeling through mechanisms including preferential activation of the ATF6 signaling arm of the unfolded protein response (UPR). However, the protein target(s) of AA263 and the potential for further development of this class of ER proteostasis regulators had not been previously explored. Here, we employ chemical proteomics to demonstrate that AA263 covalently targets a subset of ER protein disulfide isomerases, revealing a potential molecular mechanism for the activation of ATF6 afforded by this compound. We then use medicinal chemistry to establish next-generation AA263 analogs showing improved potency and efficacy for ATF6 activation, as compared to the parent compound. Finally, we show that treatment with these AA263 analogs enhances secretory pathway proteostasis to correct the pathologic protein misfolding and trafficking of both a destabilized, disease-associated α1-antitrypsin (A1AT) variant and an epilepsy-associated GABA_A_ receptor variant. These results establish AA263 analogs with enhanced potential for correcting imbalanced ER proteostasis associated with etiologically-diverse protein misfolding disorders.

## INTRODUCTION

Nearly one-third of the human proteome is targeted to the endoplasmic reticulum (ER) for folding and trafficking to downstream secretory environments including the plasma membrane, lysosome, and the extracellular space.^1–3^ The folding and trafficking versus degradation of these secretory proteins is decided by a process termed ER quality control, wherein ER-localized chaperones and folding factors engage proteins within the ER to facilitate their folding into a folded, trafficking-competent conformation.^1,2,4^ Proteins unable to attain a folded conformation in the ER through interactions with ER chaperones are instead recognized by ER-localized degradation factors that promote their targeting to degradation pathways such as ER-associated degradation by the proteasome or ER-phagy accomplished by the lysosome.^4^ Through this ER quality control process, cells promote the trafficking of folded, functional proteins through the secretory pathway and prevent the accumulation of non-native or aggregation-prone conformations within the ER or in downstream secretory environments.^5,6^

Despite the efficacy of ER quality control, failure of this process is implicated in the onset and pathogenesis of numerous, etiologically-diverse diseases, collectively referred to as protein misfolding diseases.^7^ These diseases are primarily associated with destabilizing mutations within a secretory protein that hinder ER quality control machinery, ultimately leading to toxic protein aggregation and/or loss of function pathologies. For example, the aberrant secretion and toxic extracellular aggregation of destabilized, aggregation- prone variants of amyloidogenic proteins including transthyretin (TTR) or immunoglobulin light chain (LC) are implicated in the onset and pathogenesis of protein aggregation diseases such as the TTR amyloidoses and Light Chain Amyloidosis.^8,9^ Alternatively, mutations in subunits of the gamma-aminobutyric acid type A (GABA_A_) receptors prevent the proper folding, assembly and trafficking of these proteins to the plasma membrane. This loss-of-function phenotype has been implicated in numerous neurological disorders, including genetic epilepsy and neurodevelopmental delay.^10,11^ Other proteinopathies, such as alpha-1-antitrypsin (AAT) deficiency can involve both toxic protein aggregation and loss of function pathologies. In this disease, destabilized AAT variants (e.g., PIZZ) can lead to both gain of toxic function pathology through intra-ER retention causing liver pathology and loss of function pathology associated with poorly functional secreted AAT leading to chronic obstructive pulmonary disease (COPD) in the lung.^12,13^

The central involvement of ER quality control in the onset and pathogenesis of protein misfolding diseases has suggested that developing strategies to enhance ER quality control offers a potential opportunity to broadly mitigate pathologies linked to the aberrant folding or degradation of proteins targeted to the ER.^14^ One attractive strategy to regulate ER quality control is through targeting the unfolded protein response (UPR) – the predominant stress responsive signaling pathway that regulates ER quality control in response to pathologic ER insults.^15–17^ The UPR comprises three integrated signaling pathways activated downstream of the ER membrane proteins IRE1, PERK and ATF6.^18^ ER stress activates the UPR leading to downstream transcriptional and translational signaling that functions to both alleviate the ER stress and enhance ER function. ER quality control is primarily regulated by the UPR-associated transcription factors XBP1s (downstream of IRE1) and ATF6 (a cleaved product of full length ATF6).^19,20^ These transcription factors induce expression of overlapping, but distinct, sets of ER chaperones, folding enzymes, and degradation factors that remodel the composition and the capacity of ER quality control pathways by creating emergent functions.^21,22^ Consistent with this, stress-independent activation of XBP1s, or especially, ATF6 has been shown to correct pathologic imbalances in ER quality control implicated in the pathogenesis of many different protein misfolding diseases, including those indicated above.^23–25^

The potential to improve ER quality control in protein misfolding diseases through activation of ATF6 led to a significant interest in developing pharmacologic approaches that selectively induce activation of this UPR transcriptional program. Towards that aim, we previously used high throughput screening to identify first-in-class ER proteostasis regulator compounds, such as compound AA147, that preferentially activate the ATF6 signaling arm of the UPR.^26^ We showed that AA147 activates ATF6 through a mechanism involving metabolic conversion of the prodrug AA147 to an electrophilic quinone methide on the ER membrane that covalently modifies a subset of ER protein disulfide isomerases (PDIs).^27^ This leads to an increase in reduced ATF6 that can traffic to the Golgi for proteolytic release of the active ATF6 transcription factor domain.^28–31^ Intriguingly, treatment with AA147 improves ER quality control for many different destabilized, disease-associated proteins linked to protein aggregation and loss-of-function protein misfolding diseases including AAT deficiency and GABA_A_ receptor- associated epilepsy, as well as protein aggregation-associated degenerative diseases such as TTR amyloidosis and immunoglobulin light chain amyloidosis.^26,32–34^ As predicted based on the mechanism of action of AA147, these benefits can be attributed to ER proteostasis remodeling induced by either direct PDI modification and/or by ATF6 activation, demonstrating the broad potential for this approach to influence ER proteostasis of multiple different disease-relevant proteins.^35,36^ However, despite this promise, the continued development of AA147 has proven challenging, as AA147 has been shown to be largely recalcitrant towards scaffold modifications aimed at improving its potency, selectivity, and efficacy for ATF6 activation.^27,37^

In addition to finding AA147, our original high-throughput screen also identified the phenylhydrazone compound AA263 as a compound that preferentially activates the ATF6 arm of the UPR.^26^ However, the mechanism of AA263-dependent ATF6 activation and the potential for the development of improved analogs based on this scaffold by medicinal chemistry have not been previously explored. Herein, we show that AA263, like AA147, covalently targets ER PDIs, providing a potential mechanism to explain the preferential ATF6 activation afforded by this compound. Intriguingly, we also show that modification of the AA263 B-ring at the *para* position allows for the establishment of compounds with improved selectivity and efficacy for ATF6 activation. Taking advantage of this, we establish next generation AA263 analogs that show improved potency and efficacy in comparison to the parent AA263. Further, we demonstrate that these enhanced AA263 analogs correct ER quality control defects in cellular models expressing disease-relevant proteins including the Z-variant of AAT and epilepsy-associated variants of GABA_A_ receptors. Collectively, these results establish AA263 and its associated analogs as ER proteostasis regulators with enhanced potential for further development, expanding the toolbox of compounds suitable for targeting ER quality control in the context of etiologically diverse protein misfolding diseases.

## RESULTS

### AA263 covalently modifies a subset of ER PDIs

ER proteostasis regulators including AA147 and AA132 activate the ATF6 arm of the UPR through a mechanism involving their metabolic activation to a quinone methide and subsequent covalent targeting of ER PDIs.^27,38^ Intriguingly, 2-hydroxyphenylhydrazones are known to tautomerize to a quinone methide (AA263-QM), suggesting that AA263 could similarly activate ATF6 through spontaneous quinone methide electrophile generation and covalent PDI targeting (**Fig. 1A**), without the requirement of metabolic activation.^39^ Interestingly, co-treatment with β-mercaptoethanol (BME), which acts as both an exogenous nucleophile and modulator of cellular redox potential, or the antioxidant resveratrol decreased AA263-dependent activation of the ERSE-FLuc reporter (**Fig. 1B,C**) suggesting that an ER enzyme oxidative activation of AA263 to an unknown electrophile could also be playing a role in PDI conjugation. Consistent with this, synthesis of an AA263 analog lacking the hydroxyl moiety on the A-ring (AA263-1; **Fig. S1A**) does not activate the ATF6-selective ERSE-FLuc reporter in HEK293 cells. These results support a model whereby AA263 activates ATF6 through a mechanism involving both enzymatic oxidation and direct tautomerization to an AA263-QM and subsequent covalent modification of predominantly ER protein targets (**Fig. 1A**).

**Figure 1:**
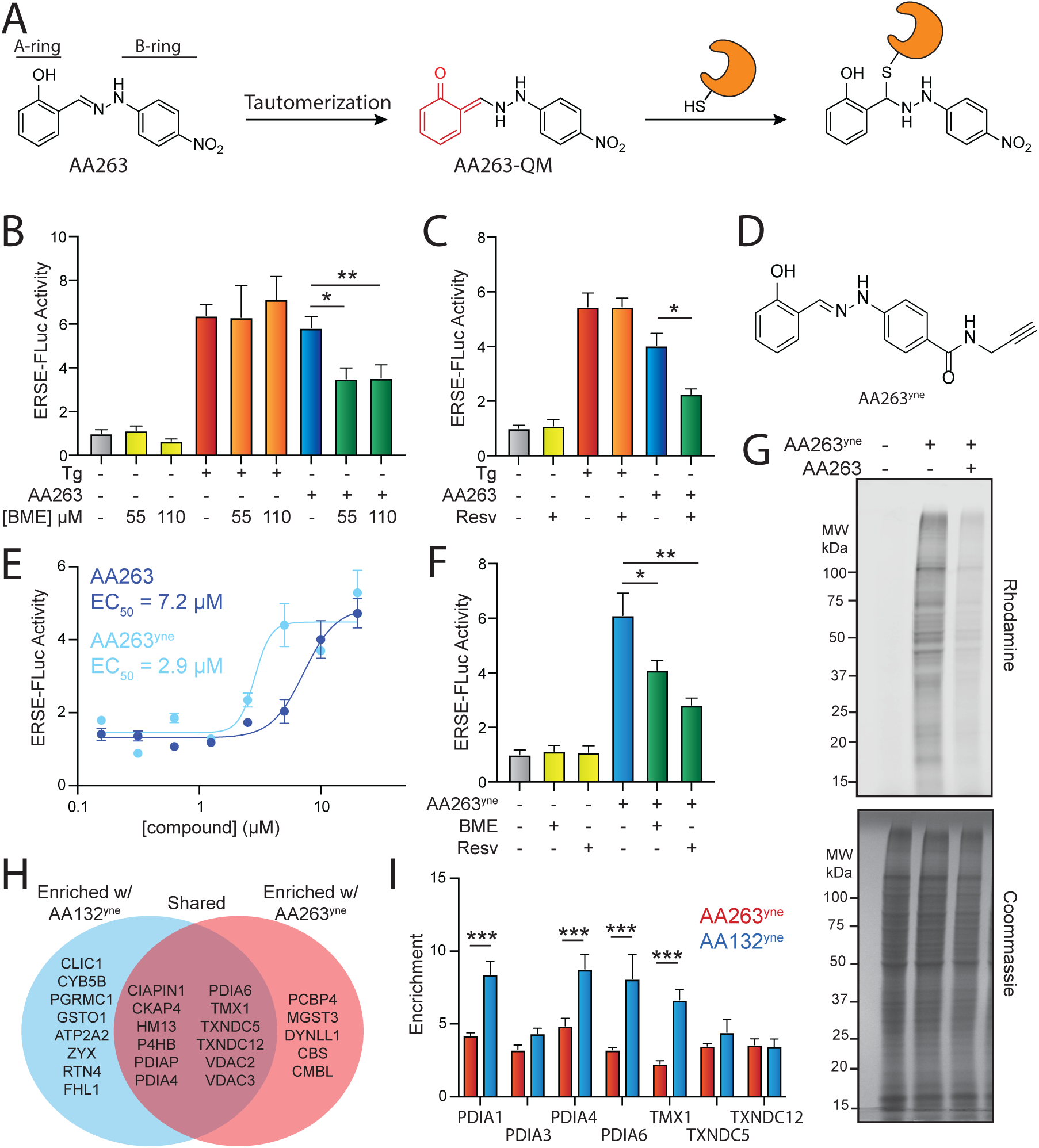
AA263 covalently modifies protein disulfide isomerase family members. **A.** Mechanism of AA263 metabolic activation and covalent protein modification **B.** Activation of the ERSE.FLuc ATF6 reporter in HEK293T cells treated for 18 hr with AA263 (10 µM) or thapsigargin (Tg, 500 nM) in the presence or absence of β-mercaptoethanol (BME; 55 μM or 110 μM). Error bars show SEM for N>6 replicates. *p<0.05, **p<0.01 for unpaired t-test. **C.** Activation of the ERSE.FLuc ATF6 reporter in HEK293T cells treated for 18 hr with AA263 (10 µM) or Tg (500 nM) in the presence or absence of resveratrol (2.5 µM). Error bars show SEM for N>6 replicates. *p <0.05 for unpaired t-test. E. **D.** Structure of AA263^yne^ **E.** Activation of the ERSE.FLuc ATF6 reporter in HEK293T cells treated for 18 h with the indicated dose of AA263 or AA263^yne^. Error bars show SEM for n=3 replicates. The EC_50_ is shown. **F.** Activation of the ERSE.FLuc ATF6 reporter in HEK293T cells treated with AA263^yne^ (10 µM) in the presence or absence of BME (55 μM) or resveratrol (2.5 μM). *p<0.05, **p<0.01 for unpaired t-test. **G.** Representative SDS-PAGE gel of Cy5-conjugated proteins from HEK293T cells treated for 4 h with vehicle (0.1% DMSO), AA263^yne^ (5 µM), or the combination of AA263^yne^ (5 µM) and AA263 (20 μM). Coomassie stained gel is shown below. **H.** Venn diagram of identified targets of AA132^yne^ and AA263^yne^. Hits defined as proteins with a significant fold change greater than 3 (p<0.01) that were identified in two independent biological experiments. **I.** TMT reporter ion enrichment ratio of select PDIs from comparative chemoproteomic experiment in HEK293T cells treated with the indicated compound relative to DMSO (n = 8 biological replicates). ***p<0.005 for a two-way ANOVA.

To monitor covalent protein labeling by AA263, we generated an AA263 analog with replacement of the nitro group on the B-ring with an alkyne amenable for affinity enrichment experiments (AA263^yne^ **Fig. 1D**). We found that this analog robustly activated the ERSE-FLuc reporter, demonstrating an ∼2-fold increase in potency, as compared to AA263 (**Fig. 1E**). Co-treatment with the selective ATF6 inhibitor Ceapin-A7 (CP7)^40–42^ blocked AA263^yne^-dependent increases in ERSE-Fluc activity, confirming that this effect can be attributed to ATF6 activation (**Fig. S1B**). We also confirmed that the AA263^yne^-dependent activation of the ERSE-FLuc reporter was inhibited by co-treatment with BME or resveratrol (**Fig. 1F**). We treated HEK293 cells with AA263^yne^ and visualized protein modification by appending a Cy5-azide fluorophore to the terminal alkyne of conjugated proteins using Cu(I)-catalyzed azide-alkyne cycloaddition (CuAAC). AA263^yne^ showed robust protein labeling in HEK293 cells upon in site compound treatment (**Fig. 1G**). Co-treatment with excess AA263 reduced labeling by AA263^yne^, confirming that AA263^yne^ and AA263 label the same subset of the human proteome. Further, co-treatment with BME or resveratrol also reduced AA263^yne^ proteome labeling (**Fig. S1C**). Intriguingly, unlike AA147^yne^, treatment of lysates with AA263^yne^ showed mild proteome labeling, indicating that some AA263-dependent protein labeling may proceed through direct tautomerization to the electrophilic QM species (**Fig. S1D**).

We next sought to identify proteins modified by AA263 using Tandem Mass Tag (TMT)-based quantitative proteomics. We previously showed that preferential ATF6 activation afforded by compounds such as AA147 and AA132 is attributed to the extent of compound-dependent modification of PDIs. ^38^ Thus, we compared the extent of proteome labeling observed with AA263^yne^ labeling to that observed with AA132^yne^ – a compound that globally activates the UPR through covalently modifying a large number of PDIs.^38^ For these comparative chemoproteomic experiments, we treated HEK293 cells with vehicle, AA263^yne^ or AA132^yne^ and then used CuAAC to append biotin-azide to modified proteins. Proteins were then enriched with streptavidin, digested with trypsin digestion, labeled with tandem mass tag (TMTs), and analyzed by LC-MS/MS.^43,44^ Modified proteins were defined by a 3-fold enrichment in compound-treated cells (*p*<0.01), as compared to vehicle-treated cells. We identified 17 proteins covalently modified by AA263^yne^, as compared to 20 proteins modified by AA132^yne^ (**Fig. 1H**). Intriguingly, 12 proteins were shared between these two conditions, including 7 different ER-localized PDIs (**Fig. 1H**). This includes PDIs previously shown to regulate ATF6 activation including TXNDC12/ERP18.^45,46^ These results are similar to those observed when comparing proteins modified by the preferential ATF6 activating compound AA147^yne^ and AA132^yne^.^38^ Further, we found that the extent of labeling for PDIs including PDIA1, PDIA4, PDIA6, and TMX1, but not TXNDC12, showed greater modification by AA132^yne^, as compared to AA263^yne^ (**Fig. 1I**). Similar results were observed for AA147^yne^.^38^ This suggests that, like AA147, the preferential activation of ATF6 afforded by AA263 is likely attributed to the modifications of a subset of multiple different ER-localized PDIs by this compound. Four of the 5 proteins that formed conjugates with AA263^yne^, but not AA132^yne^, are cytosolic proteins, suggesting that tautomerization to an AA263-QM may be occurring as a minor quinone methide formation pathway. However, spontaneous tautomerization to the quinone methide of AA263 seems unlikely to be the main pathway of electrophile formation from AA263 given the remarkable preference of AA263 to react with ER-localized proteins, which is not a general feature of other small molecule electrophiles.^47,48^ This specificity for ER proteins instead suggests the localized generation of AA263 quinone methides at the ER membrane, likely through metabolic activation by different ER localized oxidases, which has previously been shown to contribute to the selective modification of ER proteins afforded by other compounds such as AA147.^49^

### Structure activity relationship studies identify AA263 analogs with improved potency and efficacy

The increased potency observed for AA263^yne^, as compared to AA263, suggests that the B-ring of AA263 may be amenable to medicinal chemical modification for modulating the activation of ATF6 by this class of compounds (**Fig. 1E**). Further, nitro groups are known to alter mitochondrial function, so removal of this functional group could improve cellular tolerability and reduce cytotoxicity associated with two successive 1 electron reductions of the nitro group to afford cytotoxic aromatic nitroso compounds.^50,51^ Towards that aim, we synthesized a panel of AA263 derivatives comprising a heteroaromatic B ring and aryl B rings with varying functional groups largely but not exclusively appended at the para-position of the B ring (**Fig. 2A**). We screened these compounds for ATF6 activation using HEK293 cells stably expressing the ATF6-selective ERSE-Fluc reporter (**Fig. 2B**). This effort demonstrated that ATF6 activators are tolerant of B-ring *para* amide and ester functional groups, reflected by AA263^yne^ and AA263-5 being the two most active AA263 analogs. Interestingly, the *para* carboxylate substitution affords an inactive compound (AA263-4). We confirmed that AA263^yne^ and AA263-5 induced expression of the ATF6 target gene *HSPA5/BiP* in HEK293 cells (**Fig. S2A**) to levels higher than that observed for AA263, without increasing expression of the IRE1/XBP1s target gene *ERdj4* (**Fig. S2A**) or the PERK/ISR target gene *CHOP/DDIT3* (**Fig. S2A**). The inactive compound AA263-2 (**Fig. 2A**) featuring a heterocyclic B-ring was used as a control, and did not show activation of UPR target genes. ATF6 activators AA263^yne^ and AA263- 5 also both showed increased potency for ERSE-FLuc activation relative to AA263, further highlighting the improved activity of these two compounds (**Fig. S2B**)

**Figure 2.**
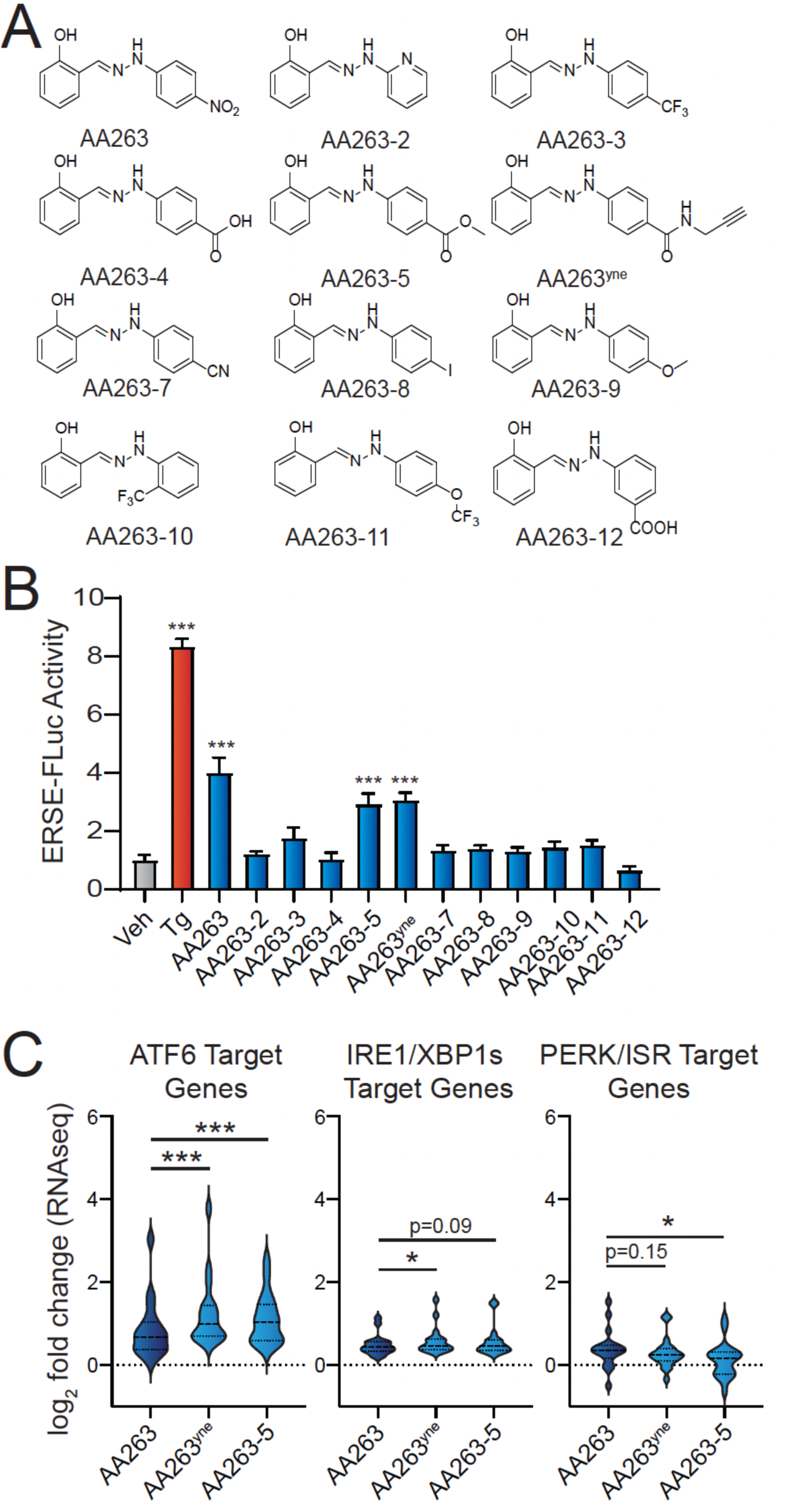
Identification of AA263 analogs that show enhanced ATF6 activation. **A.** Structures of AA263 analogs. **B.** Activation of the ERSE.Fluc ATF6 reporter in HEK293T cells reporter treated for 18 h with vehicle, Tg (0.5 μM), or the indicated analog (10 µM). Error bars show SEM for n=3-6 biological replicates. ***p<0.005 from one-way ANOVA. **C.** Expression, measured by RNAseq, of gene sets comprising target genes regulated downstream of the ATF6 (left), IRE1/XBP1s (middle), or PERK/ISR (right) arms of the UPR in HEK293T cells treated for 6 h with 10 µM AA263, AA263^yne^, or AA263-5. Full RNAseq data and genesets used in this analysis are shown in **Table S1**. *p<0.05, ***p<0.005 for one-way ANOVA.

Next, we used RNAseq to further probe the selectivity for ATF6 activation in HEK293 cells treated with either vehicle, AA263, AA263^yne^ or AA263-5. Initially, we monitored the activation of ATF6 and other arms of the UPR by comparing the expression of sets of genes regulated by these different UPR signaling pathways.^52^ Aligning with our qPCR results, AA263^yne^ and AA263-5 increased expression of ATF6 target genes, in comparison to AA263 (**Fig. 2C**, **Table S1**). These compounds also showed a modest increase in IRE1/XBP1s target genes, reflecting the known overlap between ATF6 and IRE1/XBP1 target genesets.^22^ In contrast, both compounds modestly reduced expression of PERK/ISR target genes. This may result from removal of the nitro group, as this geneset can be induced by mitochondrial dysfunction resulting from the presence of this moiety.^53,54^ These results indicate that AA263^yne^ and AA263-5 both increase adaptive ATF6 signaling, while maintaining or improving preferential selectivity for this specific arm of the UPR over the other two UPR arms. Importantly, we did not observe activation of genesets regulated downstream of other stress-responsive signaling pathways such as the oxidative stress response (OSR) or heat shock response (HSR) in HEK293 cells treated with these other AA263 analogs (**Fig. S2C, Table S1**). Further, Gene Ontology (GO) analysis showed that AA263^yne^ and AA263- 5 primarily induced expression of genes involved in biological pathways linked to ER proteostasis and UPR activation (**Table S2**).^55^ Collectively, these results indicate that AA263^yne^ and AA263-5 both show enhanced activation of the ATF6 transcriptional program, without sacrificing the transcriptome wide preferential selectivity for ATF6 activity observed for the parent compound AA263.

We next sought to evaluate whether we could further elaborate the *para*-amide substructure of the AA263^yne^ scaffold by a medicinal chemistry strategy to provide additional enhancement of ATF6 activity. Towards that aim, we synthesized AA263^yne^ analogs where the alkyne moiety was replaced with various aliphatic or aromatic substituent groups (**Fig. 3A**). We then monitored activation of the ERSE-FLUC ATF6 reporter stably expressed in HEK293 cells treated with increasing doses of these analogs. This identified two analogs, AA263- 15 and AA263-20, that showed higher levels of reporter activation, as compared to AA263 or AA263^yne^, with exhibiting increased potency (**Fig. 3B,C**). Both AA263-15 and AA263-20 increased expression of the ATF6 target gene *BiP* at 6 h after 5 µM treatment to levels greater than that observed for AA263^yne^ in HEK293 cells (**Fig. 3D**). However, these compounds did not significantly influence expression of the IRE1/XBP1s target gene *ERDJ4* or the PERK target gene *CHOP* in these cells (**Fig. S3A**). Further, we confirmed that both AA263-15 and AA263- 20 competed with proteome labeling afforded by AA263^yne^ (**Fig. S3B**), demonstrating that these compounds covalently targeted a similar subset of the proteome. These results demonstrate the potential for further developing AA263 analogs for preferential ATF6 activation and identify two compounds, AA263-15 and AA263- 20 as compounds with improved activity relative to AA263 or AA263^yne^.

**Figure 3.**
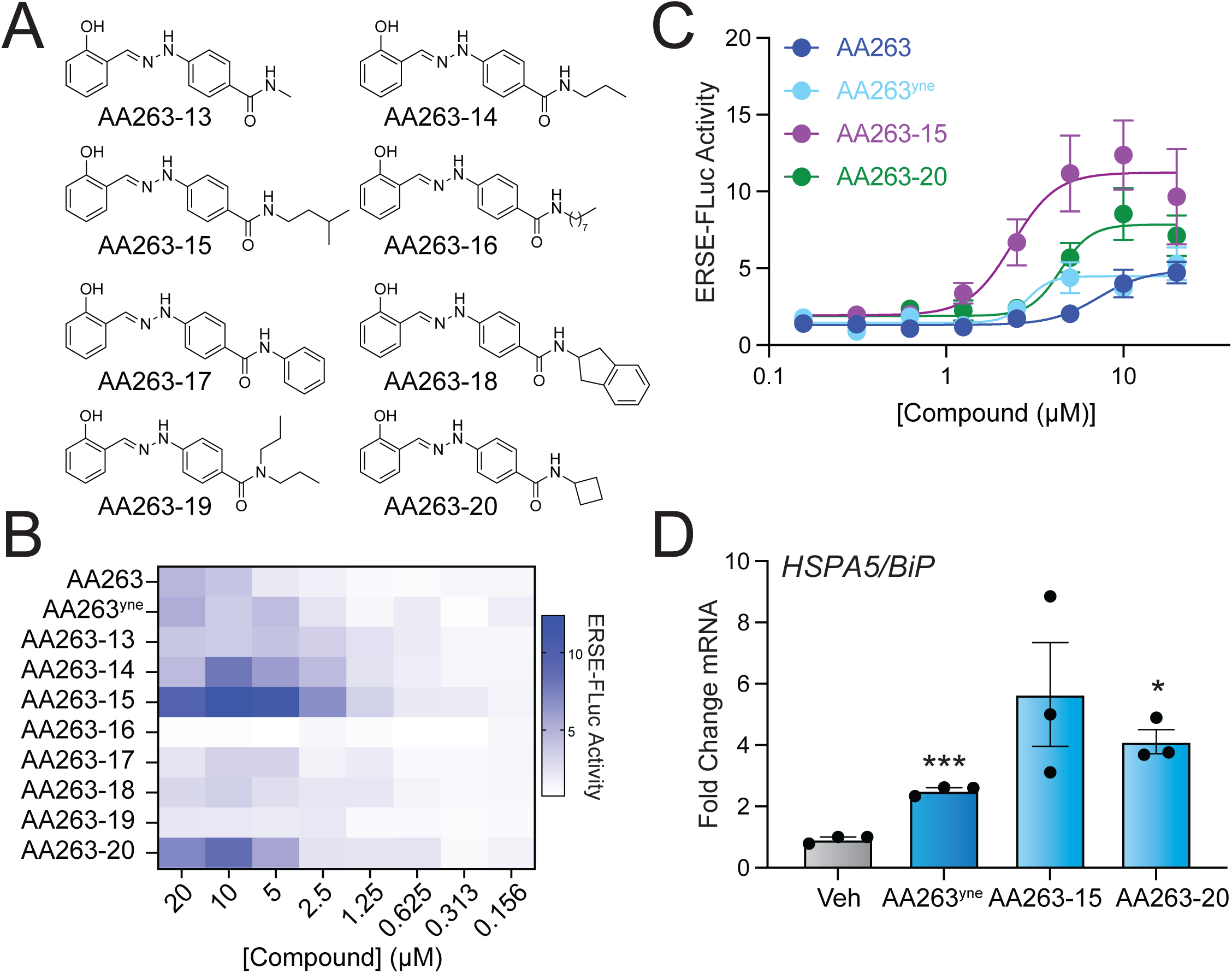
Diversification of the AA263 B-Ring affords improved AA263 analogs. **A**. Structures of AA263 analogs. **B.** Heat map showing activation of the ERSE-FLuc ATF6 reporter in HEK293T cells treated for 18 h with the indicated dose of compound. **C**. Activation of the ERSE.Fluc ATF6 reporter in HEK293T cells treated for 18 h with the indicated dose of compound. Error bars show SEM for n=6 replicates. **D.** Expression, measured by qPCR, of the ATF6 target gene *BiP* in HEK293T cells treated with indicated AA263 analog (10 µM) for 6 h. Error bars show SEM for n=3 independent biological replicates. *p<0.05, ***p<0.001 for one-way ANOVA.

### Enhanced AA263 analogs improve secretory proteostasis for the disease-associated Z-variant of A1AT

We next sought to define the potential of AA263 analogs for promoting adaptive ER remodeling in cellular models of protein misfolding diseases. AAT deficiency is a complex, multi-tissue disorder caused by the pathologic aggregation of destabilized AAT variants in the ER of hepatocytes comprising the liver and the reduced ability of secreted mutant AAT to inhibit neutrophil elastase, leading to lung dysfunction.^12^ Previously, we showed that treatment with ER proteostasis factors such as AA263 reduced intra- and extracellular AAT polymers and improved neutrophil elastase (NE) inhibitor activity of secreted AAT in HUH7.5^AAT^ ^Null^ cells transiently expressing mutant Z variant AAT .^33^ Here, we compared the potential for AA263 and improved analogs of AA263 including AA263^yne^ and AA263-20 to mitigate polymer accumulation and reduced activity of the disease-associated Z variant of AAT stably expressed in liver-derived Huh7 cells (Huh7.5Z). Initially, we confirmed that AA263, AA263^yne^, and AA263-20 induced expression of the ATF6 target gene *BiP* in these cells (**Fig. S4**). As reported previously, we found that AA263 reduced intracellular and secreted AAT-Z polymers, as measured by ELISA (**Fig. 4A,B**). However, we did not observe AA263-dependent enhancement of AAT NE inhibition activity in conditioned media, as measured by neutrophil elastase substrate-based fluorogenic assay (**Fig. 4C**).^56^ In contrast, both AA263^yne^ and AA263-20 both reduced intracellular and secreted AAT polymers and enhanced AAT-Z NE inhibition in conditioned media (**Fig. 4A-C**). This indicates that AA263^yne^ and AA263-20 both show enhanced potential to rescue AAT-Z secretory proteostasis, as compared to AA263.

**Figure 4.**
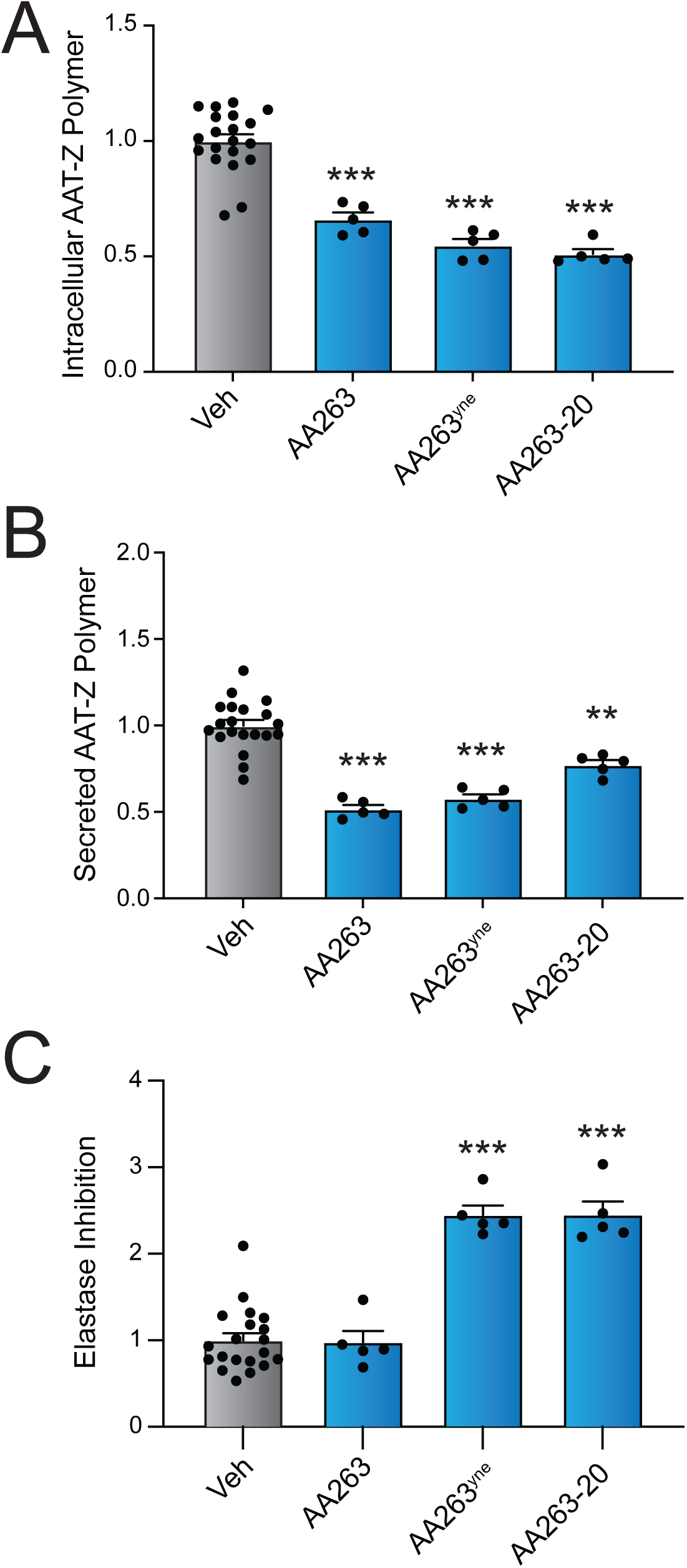
AA263 analogs improve secretory proteostasis for the disease-associated AAT-Z variant. **A-C**. Intracellular AAT-Z polymer levels (**A**), extracellular AAT-Z polymer levels in conditioned media (**B**), and elastase inhibition activity of AAT-Z in conditioned media (**C**) from Huh7.5Z cells treated for 24 h with AA263 (10 µM), AA263^yne^ (10 µM), or AA263-20 (10 µM). Error bars show SEM for n>5 replicates. Data are shown normalized to vehicle-treated cells. *p<0.05, **p<0.01, ***p<0.005 for one-way ANOVA compared to vehicle-treated cells.

### Optimized AA263 analogs rescue the surface trafficking and function of an epilepsy-prone GABA_A_ receptor variant

We next sought to investigate the potential of prioritized AA263 analogs for modulating the expression and functional activity of destabilized gamma-aminobutyric acid type A (GABA_A_) receptors. GABA_A_ receptors are pentameric proteins assembled in the ER from eight subunit classes, with the most common composition being 2 α1 subunits, 2 β2/β3 subunits, and 1 γ2 subunit.^57,58^ Various variants in these major subunits have been reported to impair the trafficking of GABA_A_ receptors to the plasma membrane, ultimately leading to a significant decrease in their function.^59,60^ As such, GABA_A_ variants have been widely associated with the onset and pathogenesis of numerous neurological disorders, including genetic epilepsy.^11,59,61^ It has previously been reported that enhancement of ER proteostasis can stabilize GABA_A_ variants and restore their surface trafficking as well as their function.^34,62^ Thus, we aimed to determine if our optimized AA263 analogs could similarly rescue the surface expression and activity of a known trafficking-deficient γ2(R177G) GABA_A_ variant, associated with complex febrile seizures.^63^

We treated HEK293T cells stably expressing α1β2γ2(R177G) pathogenic receptors with each AA263 analog. While both AA263^yne^ and AA263-5 increased total protein levels of γ2(R177G) (**Fig. S5A**), only AA263^yne^ could increase the surface expression of α1β2γ2(R177G) GABA_A_ receptors transiently transfected in HEK293T cells (**Fig. 5A**). Thus, the optimized compound AA263^yne^ effectively rescues the trafficking of this GABA_A_ variant to the cell surface. Importantly, the increased total protein levels of γ2(R177G) afforded by treatment with AA263^yne^ could be partially attenuated by co-treatment with the selective ATF6 inhibitor Ceapin-A7 (CP7) ^40–42^, indicating that compound-dependent ATF6 activation contributes to this observed increase (**Fig. S5B**). Next, we used cycloheximide (CHX) chase assays to determine the impact of AA263^yne^ on the turnover of the γ2(R177G) variant. Compared with WT receptors, we found that the pathogenic γ2(R177G) variant displayed a faster turnover, which was slowed down by treatment with AA263^yne^ to a similar rate compared to GABA_A_ WT (**Fig. S5C**). In addition to AA263^yne^, both AA263-15 and AA263-20 increased the total expression levels of γ2(R177G) subunit in HEK293T cells treated with either compound (**Fig. S5D**). Further, treatment with these compounds increased surface expression levels of the γ2(R177G) subunit (**Fig. 5B**). Together, these results show that AA263^yne^ and optimized analogs such as AA263-20 can efficiently correct the misfolding of an epilepsy- associated GABA_A_ variant and rescue its trafficking to the plasma membrane.

**Figure 5.**
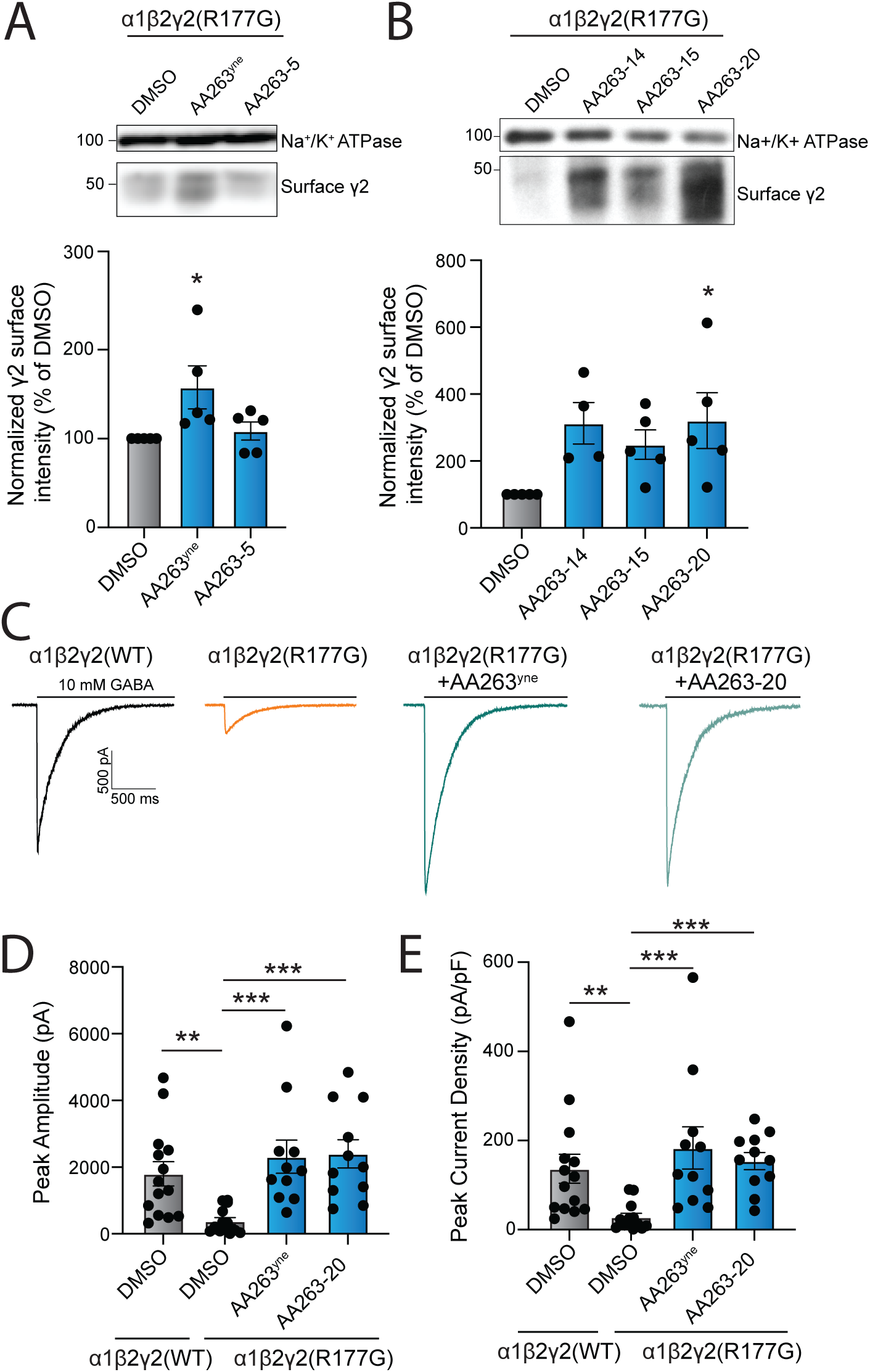
Enhanced AA263 analogs promote the trafficking and plasma membrane activity of destabilized, disease-associated GABA_A_ receptors. **A.** Representative immunoblot (above) and quantification (below) of surface biotinylated γ2 in HEK293T cells stably expressing α1β2γ2(R177G) GABA_A_ receptors treated for 24 h with 10 µM AA263^yne^ or AA263-5. Na^+^/K^+^ ATPase serves as a loading control. **B.** Representative immunoblot showing surface γ2 expression in HEK293T cells transiently transfected with α1β2γ2(R177G) receptors and treated with indicated AA263 analogs (10 µM, 24hrs. Na^+^/K^+^ ATPase serves as a loading control. **C**. Representative eIPSC traces for α1β2γ2(WT) GABA_A_ receptors and, α1β2γ2(R177G) GABA_A_ receptors treated for 24 h with DMSO, AA263^yne^ (10 µM), or AA263-20 (10 µM). 10 mM GABA (saturating condition) was applied to the recorded cells to evoke currents. **D,E**. Histograms showing changes in eIPSC peak amplitude (D) and peak current density (E) for the indicated groups. Band intensities were quantified using ImageJ software, normalized to the DMSO control condition. Each data point is reported as mean ± SEM. One-way ANOVA followed by post-hoc Dunnett’s test was used for statistical analysis for A and B. Kruskal-Wallis test followed by post-hoc Dunn’s test was used for statistical analysis for D and E. *p<0.05; **p<0.01; ***p<0.001.

Finally, to determine if compound-dependent increases of α1β2γ2(R177G) receptor surface expression also translate into a rescued function of the receptor, we used whole-cell patch clamp in HEK293T cells transiently transfected with either WT GABA_A_ receptor or the α1β2γ2(R177G) GABA_A_ variant and recorded evoked inhibitory postsynaptic currents (eIPSCs) by a 4-second application of 10 mM GABA. Consistent with the literature we first showed that cells expressing the epilepsy-associated γ2(R177G) variant displayed eIPSCs of significantly smaller amplitude than cells expressing WT GABA_A_ receptors (**Fig 5C,D**).^64^ In addition, after normalizing each cell’s amplitude by its capacitance, γ2(R177G) GABA_A_-containing cells had a lower peak current density when compared with WT GABA_A_-containing cells (**Fig. 5C,E**). Importantly, treatment with AA263^yne^ or AA263-20 in HEK293T cells transiently expressing α1β2γ2(R177G) GABA_A_ receptors restored both eIPSC peak amplitude and peak current density to levels nearly identical to those observed in cells expressing WT GABA_A_ receptors (**Fig. 5C-E**). Collectively, these findings establish AA263^yne^ and AA263-20 as ER proteostasis regulators that not only restore surface trafficking of disease-associated GABA_A_ receptor variants, but also rescue their functional activity at the cell surface.

## DISCUSSION

Herein, we define the molecular basis for AA263-dependent ER proteostasis remodeling. We show that AA263 covalently targets a subset of ER-localized PDIs, suggesting that this compound, like AA147, promotes activation of the adaptive ATF6 signaling arm of the UPR through a mechanism involving PDI modification. Intriguingly, unlike AA147, structure-activity relationship studies showed that AA263 activity could be enhanced by altering the putative B-ring of this compound. This identified two analogs, AA263-5 and AA263^yne^, that exhibited enhanced selectivity and efficacy for ATF6 activation. Further optimization of the amide moiety of AA263^yne^ afforded prioritized analogs including AA263-20. We then showed AA263^yne^ and this prioritized analog exhibited enhanced activity relative to the parent compound AA263 in correcting ER proteostasis in cellular models of alpha-1- antitrypsin deficiency and GABA_A_ receptor channelopathies. These studies demonstrate the potential to enhance ER proteostasis remodeling activity with compound AA263 to correct proteostasis deficits in etiologically-diverse protein misfolding diseases.

Unlike other ER proteostasis activators, such as AA147 and AA132, AA263 does not possess the 2- amino-*p*-cresol moiety required for the enzymatic activation of these other compounds to a reactive quinone methide. Despite this, we show that AA263 functions through the covalent targeting of ER-localized PDIs. Salicaldehyde N-acylhydrazones can generate transient quinone-methide type species driven by tautomerization.^65^ This suggests that AA263 tautomerization to a quinone methide may underlie this covalent PDI targeting. Consistent with this, we show that AA263 can covalently modify proteins ex vivo in cellular lysates. However, a key question is how does AA263 show selectivity for ER PDIs independent of the localized metabolic activation afforded by other compounds such as AA147. One potential explanation is that the AA263 electrophilic species may possess differential reactivity towards a subset of proteinogenic thiols such as those in PDIs versus glutathione.^66,67^ Regardless, these results, when combined with our analyses for AA147 and AA132^49^, demonstrate how the diverse covalent engagement profiles of subtly different electrophilic species against a subset of ER proteins can elicit a spectrum of transcriptional outcomes.

Our findings that AA263 engages similar PDI family members and ER proteins as AA147 further suggesting the importance of this family of proteins for adjusting ER proteostasis through mechanisms including activation of ER stress responsive signaling pathways.^46,68–70^ We previously showed that the extent of PDI labeling directly impacts the ability for this class of compounds to activate ATF6, with more modest levels leading preferential ATF6 activation while higher levels of labeling activating all three arms of the UPR.^49^ Interestingly, our results indicate that AA263 maintains preferential selectivity for ATF6 activation despite modifying an intermediate population of PDIs greater than those observed for the selective ATF6 activator AA147, but less than those observed for the global UPR activator AA132. This suggests that higher levels of PDI engagement than those observed with AA147 are still accessible without compromising the ability for these compounds to preferentially activate ATF6.

Intriguingly, like AA147 and AA132, we show that the AA263 scaffold is amenable to modifications on the putative non-reactive B-ring. However, unlike these other compounds, medicinal chemistry efforts focused on improving AA263 activity identified analogs with improved potency and transcriptional selectivity for ATF6 activation relative to the parent compound. The improvements in transcriptional selectivity of the compounds may be attributed to the removal of the nitro group from the parent compound AA263. Nitroarene-containing compounds have been shown to affect mitochondrial function and also lead to ROS production through nitroarene metabolism.^71,72^ Thus, the crosstalk between ISR signaling and the mitochondria may explain the higher levels of PERK/ISR signaling with AA263 relative to our prioritized analogs.^73–75^ Additionally, we do observe some basal CHOP signaling with inactive AA263 analogs, exemplified by AA263-2, perhaps suggesting some inherent promiscuity of the phenylhydrazone scaffold. As a result, the identity of the B-ring electron- withdrawing group may improve the spectrum of target protein engagement in a positive manner to boost selectivity for ATF6 activation.^66^

It is important to note that both 2-hydroxyphenylhydrazones and 4-hydroxyphenylhydrazones have been classified as pan-assay interference compounds (PAINS), due, in part, to their reactivity towards biological nucleophiles and metal chelation properties.^76,77^ ^76,78^ However, our observation of SAR trends connecting scaffold modifications to magnitude of pathway activation indicates that this particular compound activity is perhaps independent of PAINS activity. Further, many of our negative analogs possess a functional A-ring, and thus would still be expected to possess covalent reactivity if PAINS behavior mediated the activation of stress-responsive signaling.^79^ Thus any nonspecific covalent modification, if it exists, seems to be functionally silent in this assay. Moreover, we found that these compounds showed remarkable specificity for ER-localized protein engagement, further indicating that AA263 and prioritized analogs possess other potential properties that drive specificity towards the ER, which we are continuing to pursue.

Optimized AA263 analogs identified herein have enhanced potential to correct ER proteostasis in two distinct cellular models of protein misfolding diseases – AAT deficiency and GABA_A_ receptor trafficking. We show that AA263 analogs reduce both intracellular and extracellular accumulation of AAT-Z aggregates and improve activity of the secreted protein, correcting both aspects of disease pathology. Further, we demonstrate that our optimized AA263 analogs enhanced trafficking, surface expression, and membrane activity of a pathogenic GABA_A_ receptor variant associated with genetic epilepsy, restoring is functional activity at the plasma membrane. AA263 and its related analogs can influence ER proteostasis in these models through different mechanisms including ATF6-dependent remodeling of ER proteostasis and direct alterations to the activity of specific PDIs. Consistent with this, we show that pharmacologic inhibition of ATF6 only partially blocks increases of γ2(R177G) afforded by treatment with AA263^yne^, highlighting the benefit for targeting multiple aspects of ER proteostasis to enhance ER proteostasis of this disease-relevant GABA_A_ variant. While additional studies are required to further deconvolute the relative contributions of these two mechanisms on the protection afforded by our optimized compounds, our results demonstrate the potential for these compounds to enhance ER proteostasis in the context of different protein misfolding diseases. As we and others continue expanding on this class of ER proteostasis regulators, we will further demonstrate the therapeutic utility of these compounds in cellular and in vivo models and identify next-generation compounds with improved potential for translation to correct ER proteostasis defects in etiologically-diverse diseases.

## MATERIALS AND METHODS

### Reagents & Plasmids

Cycloheximide (cat #01810) and γ-aminobutyric acid (#A2129) were obtained from Sigma Aldrich. Human neutrophil elastase (Cat # IHUELASD100UG) was obtained from Innovation Research. MG-132 (#A2585) was obtained from ApexBio. Thapsigargin was purchased from Sigma-Aldrich. AA263 analogs and AA263^yne^ analogs were synthesized as described in *Supplemental Materials and Methods* and used as 10 mM DMSO stocks. The pCMV6 plasmids containing human GABA_A_ receptor α1 (Uniprot no. P14867-1), β2 (isoform 2, Uniprot no. P47870-1), γ2 (isoform 2, Uniprot no. P18507-2) subunits, and pCMV6 Entry Vector plasmid (pCMV6-EV) were obtained from Origene. The human GABA_A_ receptor γ2 subunit missense mutation R177G was constructed using QuikChange II site-directed mutagenesis Kit (Agilent Genomics). All cDNA sequences were confirmed by DNA sequencing.

### Antibodies

The rabbit anti-GABA_A_R-γ2 polyclonal antibody (#AB5559) was obtained from Millipore. The rabbit monoclonal anti-Na^+^/K^+^-ATPase (#ab76020) antibody, was obtained from Abcam. The rhodamine anti-actin primary antibody (#12004163) was obtained from BioRad. Mouse anti-human AAT monoclonal antibody 2C1 was obtained from Hycult Biotech (Cat #HM2289). Mouse anti-human AAT monomer-specific monoclonal antibody 16F8 was a kind gift from the Balch Lab. The secondary goat anti rabbit antibody (#A27036) and goat anti mouse antibody (#A28177) used for western blot were obtained from Invitrogen. The secondary goat anti-mouse HRP antibody used for the AAT Elisa experiment was obtained from Thermo-Fischer scientific (Cat # 32230).

### Cell Culture

HEK293T-Rex (ATCC), HEK293T (ATCC), and MEF (ATCC) cells were cultured in high-glucose Dulbecco’s Modified Eagle’s Medium (DMEM) supplemented with glutamine, penicillin/streptomycin and 10% fetal bovine serum (FBS). Cells were routinely tested for mycoplasma every 6 months. We did not further authenticate the cell lines. All cells were cultured under typical tissue culture conditions (37 °C, 5% CO2).

### Generation of Stable HEK293T α1β2γ2(R177G) Cell Line

HEK293T cells were grown in 10cm dishes and allowed to reach ∼70% confluency before transient transfection using TransIT-2020 (Mirus Bio #MIR 5406, Madison, WI) according to the manufacturer’s instruction. Stable cell lines for α1β2γ2(R177G) were generated using the antibiotic G-418 selection method. Briefly, cells were transfected with α1β2γ2(R177G) (0.25μg:0.25μg:0.5μg) plasmids, and then selected with 0.8 mg/mL G-418 (Fisher # 50-153-2785) for 7 days. Cells were diluted and passed to 96-well plates to ensure single cell distribution per well. Cells expanded from monoclonal cells were assessed for the expression of α1, β2, and γ2 by western blot analysis. Positive monoclonal cells were selected for experimentations.

### Measurement of UPR activity using luciferase reporters

HEK293T-Rex cells expressing the ERSE.FLuc^80^, XBP1s.RLuc ^81^, or ATF4.FLuc reporter were plated at 80 µL/well from suspensions of 250,000 cells/mL in white clear-bottom 96-well plates (Corning) and incubated at 37°C overnight. The following day, cells were treated with 20 µL of compound-containing media to give final concentration as described before incubating for 18 hr at 37°C. The plates were equilibrated to room temperature, then either 100 μL of Firefly luciferase assay reagent-1 (ERSE.FLuc and ATF4.FLuc) or Renilla luciferase assay reagent-1 (XBP1s.RLuc) (Targeting Systems) were added to each well. Samples were dark adapted for 10 min to stabilize signals. Luminescence was then measured in an Infinite F200 PRO plate reader (Tecan) and corrected for background signal (integration time 500 ms). All measurements were performed in biologic triplicate.

### Quantitative RT-PCR

The relative mRNA expression levels of target genes were measured using quantitative RT-PCR. Cells were treated as described at 37°C, harvested by trypsinization, washed with Dulbecco’s phosphate-buffered saline (GIBCO), and then RNA was extracted using the QuickRNA Miniprep Kit (Zymo). qPCR reactions were performed on cDNA prepared from 500 ng of total cellular RNA using the High-Capacity cDNA Reverse Transcription Kit (Applied Biosystems). PowerSYBR Green PCR Master Mix (Applied Biosystems), cDNA, and appropriate primers purchased from Integrated DNA Technologies (see Table below) were used for amplifications (6 min at 95°C, then 45 cycles of 10 s at 95°C, 30 s at 60°C) in an ABI 7900HT Fast Real Time PCR machine. Primer integrity was assessed by a thermal melt to confirm homogeneity and the absence of primer dimers.

Transcripts were normalized to the housekeeping genes RPLP2 and all measurements were performed in biological triplicate. Data were analyzed using the RQ Manager and DataAssist 2.0 software (ABI). qPCR data are reported as mean ± standard deviation plotted using Prism GraphPad.

### Sequences of Primers for qPCR

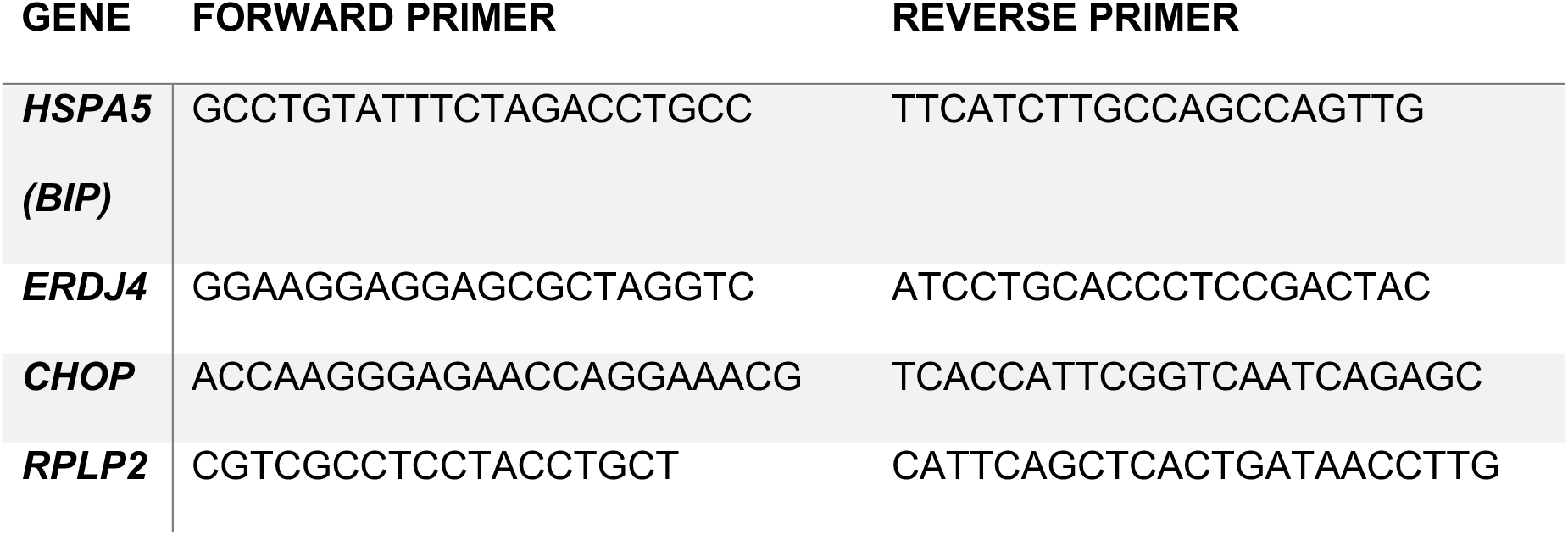

### SDS-PAGE In-Gel Fluorescence Scanning

Indicated cell line (250,000 cells/well) was treated with indicated compound or combination of compounds in six- well plates at 10 μM for six hours. Cells were lysed in radioimmunoprecipitation assay (RIPA) buffer (150 mM NaCl, 50 mM Tris pH 7.5, 1% Triton X-100, 0.5% sodium deoxycholate, and 0.1% SDS) supplemented with fresh protease inhibitor cocktail (Roche, Indianapolis, IN) and centrifuged for 10 min at 16000×*g* following a 30-minute incubation. Protein concentration of supernatant determined by BCA assay (Thermo Fisher) and normalized to give 42.5 µL at 2.35 mg/mL (100 µg/total protein). 7.5 µL ‘click chemistry master mix’ was added to each sample to give final concentrations of 100 µM of Cy5-azide (Click Chemistry Tools, Scottsdale, AZ), 800 µM copper (II) sulfate, 1.6 mM BTTAA ligand (2-(4-((bis((1-tert-butyl-1H-1,2,3-triazol-4-yl)methyl)amino)methyl)-1H-1,2,3- triazol-1-yl)acetic acid) (Albert Einstein College), and 5 mM sodium ascorbate. Reaction incubated at 30° C for 1 hour while shaking before CHCl_3_/MeOH protein precipitation. Dried protein was redissolved in 20 µL 1X SDS loading buffer and 25 µg was loaded on gel for SDS-PAGE in-gel fluorescence scanning and subsequent visualization using an Odyssey Infrared Imaging System (Li-Cor Biosciences) or BioRad gel imager.

### Comparative AA132 vs. AA263 Chemoproteomic Experiments

HEK293T cells in 10 cm plates at 80-90% confluency were treated for 6 h with vehicle (0.1% DMSO), AA132yne (10 μM), or AA263-5 (10 μM) at 37 °C. The cells were washed with PBS before harvesting with trypsin, pelleting (500 g, 5 min), and washing with PBS (1 mL). Cells pellets were resuspended in radioimmunoprecipitation assay (RIPA) buffer before sonication with a probe tip sonicator for cell lysis (15 sec, 3 sec on/2 off, 30% amplitude). For each sample, 1 g lysate (500 µL) were reacted with click reagents to give final concentrations as follows: 100 µM of diazo biotin-azide (Click Chemistry Tools, Scottsdale, AZ), 800 µM copper (II) sulfate, 1.6 mM BTTAA ligand (2-(4-((bis((1-tert-butyl-1H-1,2,3-triazol-4-yl)methyl)amino)methyl)-1H-1,2,3-triazol-1-yl)acetic acid) (Albert Einstein College), and 5 mM sodium ascorbate. The reaction was placed on a shaker at 1000 rpm at 30 °C for 90 The reaction was quenched with the sequential addition of cold methanol (4x volume), chloroform (1x volume), and DPBS (4x volume) to precipitate proteins. Proteins were pelleted by centrifugation (4,700 g, 10 min, 4 °C). The supernatant was discarded, and the pellets dried under air for 5 min. Protein pellets were resuspended in 6M urea in PBS (500 µL) with brief sonication. 50 µL of high-capacity streptavidin beads were washed with PBS and mixed with the protein solution in 6 mL of phosphate-buffered saline (PBS). This suspension was placed on a rotator or a shaker and agitated for 2 h. The beads were centrifuged and washed 5 times with PBS and 1% SDS. The protein was eluted from the beads by two washes of 50 mM sodium dithionite in 1% SDS for 1 h and then precipitated by chloroform/methanol precipitation as described above. 50 µL of freshly made 1:1 mixture 200 mM TCEP·HCl in DPBS and 600 mM K2CO3 in DPBS was added to each sample before incubation at 37 °C for 30 minutes while shaking. Alkylation of reduced thiols was achieved by addition of 70 µL freshly prepared 400 mM iodoacetamide in DPBS and incubation at room temperature while protected from light. The reaction was quenched by adding 130 µL of 10% SDS in DPBS and then diluted to approximately 0.2% SDS via DPBS (5.5 mL) and incubated with preequilibrated streptavidin agarose beads (3x1 mL PBS wash). The samples were rotated at room temperature for 1.5 hours, centrifuged at 2000 rpm for 2 minutes, and then washed sequentially with 5 mL 0.2% SDS in DPBS, 5 mL DPBS, and 5 mL 100 mM TEAB (Thermo Cat #90114) pH 8.5 to remove non-binding proteins. The beads were transferred to low-bind 1.5 mL Eppendorf tubes and the bound proteins digested overnight at 37 °C in 200 µL 100 mM TEAB containing 2 µg sequencing grade porcine trypsin, 1 mM CaCl2, and 0.01% ProteaseMax (Promega Cat #V2071). The beads were centrifuged at 2000 rpm for 5 minutes to separate the beads from the supernatant. 200 µL supernatant was transferred to a new tube using a gel-loading tip, and the beads were washed with 100 µL TEAB buffer. The beads were centrifuged at 2000 rpm for 5 minutes and the supernatant combined with the previous. 120 µL acetonitrile was added to each supernatant sample before addition of 80 µL (200 µg) of TMT 10 plex (Thermo Scientific, cat #90110) reconstituted in acetonitrile. The samples were incubated at room temperature for 1 hour and vortexed occasionally. 7 µL of freshly prepared 5% hydroxylamine in water was added to each sample to quench the reaction, vortexed, and incubated for 15 minutes before quenching with addition of 5 µL MS-grade formic acid. The samples were then vacuum centrifuged to dryness. The samples were combined by redissolving the contents of one tube in 200 µL 0.1 % trifluoroacetic acid solution in water and sequential transfer to the respective multiplexed experiment until all samples were redissolved. This stepwise process was repeated with an additional 100 µL 0.1 % TFA solution for a final volume of 300 µL. The pooled samples were fractionated using the Pierce high pH Reversed-Phase Fractionation Kit (Thermo Fisher Scientific 84868) according to manufacturer’s instructions. The peptide fractions were eluted from the spin column with solutions of 0.1% triethylamine containing an increasing concentration of MeCN (5 - 95% MeCN; 8 fractions). Samples were dried via vacuum centrifugation, reconstituted in 50 µL 0.1% formic acid, and stored at -80 °C until ready for mass spectrometry analysis.

LC-MS/MS analysis was performed using a Q Exactive mass spectrometer equipped with an EASY nLC 1000 (Thermo Fisher). The digest was injected directly onto a 30 cm, 75 µm ID column packed with BEH 1.7 µm C18 resin (Waters). Samples were separated at a flow rate of 200 nl/min on a nLC 1000 (Thermo). Buffer A and B were 0.1% formic acid in water and acetonitrile, respectively. A gradient of 5–40% B over 110 min, an increase to 50% B over 10 min, an increase to 90% B over another 10 min and held at 90% B for a final 10 min of washing was used for 140 min total run time. The column was re-equilibrated with 20 µl of buffer A prior to the injection of sample. Peptides were eluted directly from the tip of the column and nanosprayed directly into the mass spectrometer by application of 2.5 kV voltage at the back of the column. The Q Exactive was operated in a data-dependent mode. Eluted peptides were scanned from 400 to 1800 m/z with a resolution of 30,000 and the mass spectrometer in a data dependent acquisition mode. The top ten peaks for each full scan were fragmented by HCD using a normalized collision energy of 30%, a 100 ms activation time, a resolution of 7500, and scanned from 100 to 1800 m/z. Dynamic exclusion parameters were 1 repeat count, 30 ms repeat duration, 500 exclusion list size, 120 s exclusion duration, and exclusion width between 0.51 and 1.51. Peptide identification and protein quantification was performed using the Integrated Proteomics Pipeline Suite (IP2, Integrated Proteomics Applications, Inc., San Diego, CA) as described previously.

### RNA-seq

Cells were lysed and total RNA collected using the Quick-RNA Miniprep kit from Zymo Research (R1055) according to manufacturer’s instructions. RNA concentration was then quantified by NanoDrop. Whole transcriptome RNA was then prepared and sequenced by BGI Americas on the BGI Proprietary platform, which provided paired-end 50 bp reads at 20 million reads per sample. Each condition was performed in triplicate. RNAseq reads were aligned using DNAstar Lasergene SeqManPro to the Homo_sapiens-GRCh38.p7 human genome reference assembly, and assembly data were imported into ArrayStar 12.2 with QSeq (DNAStar Inc.) to quantify the gene expression levels and normalization to reads per kilobase per million. Differential expression analysis was assessed using DESeq2 in R, which also calculated statistical significance calculations of treated cells compared to vehicle-treated cells using a standard negative binomial fit of the reads per kilobase per million data to generate fold-change quantifications. The complete RNAseq data is deposited in gene expression omnibus (GEO) as GSE309691.

### Immunoblotting

Cells were grown in 6-well plates or 10-cm dishes and allowed to reach ∼70% confluency before transient transfection with α1:β2:γ2(WT) or α1:β2:γ2(R177G) (0.25μg:0.25μg:0.5μg) plasmids using TransIT-2020 (# Mirus Bio, Madison, WI) according to the manufacturer’s instruction. The mixture was incubated for 15-20 minutes at room temperature before being added to the cells. After indicated treatment, cells were harvested with Trypsin and lysed with pH 7.5 lysis buffer: (50 mM Tris, 150 mM NaCl, 2 mM *n*-Dodecyl-b-D-maltoside (DDM) (GoldBio, catalog #: DDM5)) (pH 7.5) supplemented with complete protease inhibitor cocktail (Roche #4693159001). Lysates were cleared by centrifugation (21, 000 xg, 10 min, 4°C). Protein concentrations were then measured through microBCA assay (ThermoFisher Pierce #23235). Aliquots of cell lysates were loaded with 4x Laemmli buffer (Biorad #1610747, CA) with 10% 2-Mercaptoethanol (Sigma Aldrich #M3148, Saint Louis, MO) and separated in an 8% SD-PAGE gel. The β-actin serves as a total protein loading control, whereas Na^+^/K^+^-ATPase serves as a plasma membrane protein loading control. Band intensity was quantified using Image-J software from the NIH. For both surface and total expression quantification, the protein level was first normalized to the loading control (β-actin or Na^+^/K^+^-ATPase) and then to the vehicle control (DMSO or WT).

### AAT-Z specific conformation (monomer or polymer) ELISA assay

Huh7.5Z cells stably expressing AAT-Z (E342K)^82^ were seeded in 96-well plates at a density of 2.5x10^4^ cells per well. Cells were treated with compounds for 24 h before collecting the cell culture medium. Wells were aspirated of residual media & refilled with 80uL/well of lysis buffer (50 mM Tris, 150mM NaCl, 1% (v/v) Triton X-100, and 1X Halt™ protease inhibitor cocktail) and incubated on 4°C shaker for 1 h before collecting the resultant lysate. The previous day, clear flat bottom polystyrene high binding 96-well plates were pre-coated with goat anti-human AAT polyclonal antibody (A80-122A 1:1000 in coating buffer (28.6mM g Na2CO3, 71.4mM NaHCO3, pH = 9.6)) at 4°C. Coated plates were washed 5x with PBST washing buffer (1X PBS + 0.05% (v/v) Tween-20), blocked for 2-4 h in washing buffer containing 5% dehydrated milk, and washed another 5x with to remove blocking agent just prior to collection of treated cell products. 20 μL of culture medium or lysate from each treated well was dispensed into the antibody-coated plates and incubated overnight (8-12 h) at 4°C. Plates were washed 5x with PBST washing buffer before applying conformation specific primary antibodies for AAT monomer (16F8^83^ 1:2000 in 1X PBS) or polymer (2C1^84^ 1:1000 in 1X PBS) and incubating 2-4 h at room temperature. Plates were washed 5x with PBST washing buffer before applying the secondary HRP conjugated goat-anti-mouse antibody (1:5000 in 1xPBS) and incubating 2-4 h at room temperature. After washing a final 5x with PBST washing buffer, TMB substrate (Thermo Fisher Scientific, No. 34029) was added for 10 min before quenching the reaction with 2 M sulfuric acid. Endpoint ELISA signal (450nm absorbance) was quantified by BioTek Synergy H1 Hybrid plate reader. AAT-Z monomer and polymer levels in media and lysate following 24 h treatment with AA263 analogues were normalized to vehicle treated Huh7.5Z cells.

### AAT-Z inhibitory activity (fluorogenic elastase substrate turnover) assay

Culture medium from treated Huh7.5Z cells was collected and added to pre-coated high binding 96-well plates as described above for overnight (8-12 h) incubation at 4°C. Plates were washed 5x with PBST washing buffer before dispensing a solution of porcine elastase (PPE, Promega, No. V1891) at 4 ng/well in reaction buffer (50mM Tris, pH 8.5) and incubating the plate for 1hr at 37°C. Then, 25 pmol/well of the fluorogenic elastase substrate (Z-Ala4)2Rh110 (Cayman Chemical, No. 11675) was added, and the plate incubated for 1.5 h at 37°C. The plate was read by a BioTek Synergy H1 Hybrid plate reader at 485nm excitation and 525nm emission. Elastase inhibitory activity of secreted AAT-Z in media following 24 h treatment with AA263 analogues was normalized to vehicle treated Huh7.5Z cells.

### Biotinylation of Cell Surface Proteins

HEK293T cells were plated in 6cm dishes for surface biotinylation assays. Cells were transfected with GABA_A_ receptor variants as indicated 48h prior to harvest. 24h post incubation with the indicated compounds, intact cells were washed with ice-cold Dulbecco’s Phosphate Buffered Saline (DPBS, VWR #10128-844) and incubated with the membrane-impermeable biotinylation reagent Sulfo-NHS SS-Biotin (1mg/mL; ApexBIO, #A8005) in DPBS containing 0.1mM CaCl_2_ and 1mM MgCl_2_ (DPBS + CM) for 30 min at 4°C to label surface membrane proteins. Cells were then incubated with 10mM Glycine twice at 4°C for 5 min to quench reaction. To block sulfhydryl groups, the cells were then incubated with 5nM N-ethyl-maleimide (NEM) in DPBS for 15 min at room temperature. Cells were then solubilized overnight at 4°C in lysis buffer containing 2mM DDM, 50mM Tris-HCL, 150mM NaCl, and 5mM ETA (pH 7.5) supplemented with Roche complete protease inhibitor cocktail (Roche #04,693,116,001) and 5mM NEM. To pellet cellular debris, the lysates were cleared by centrifugation (21,000 x g, 10 min, 4°C). The supernatant containing the biotinylated surface proteins was affinity-purified by incubating for 1h at 4°C with 40μL of immobilized neutravidin-conjugated agarose bead slurry (VWR #PI29201). The samples were then subjected to centrifugation (21,000 x g, 10 min, 4°C). The beads were washed three times with a buffer containing in mM: 50 Tris-HCL, 150 NaCL, and 5 EDTA (pH 7.5. Surface proteins were finally eluted from beads by vortexing for 30 min with 80μL of a buffer containing 2 x Laemmli sample buffer, 100 mM DTT, and 6 M urea (pH 6.8) for SDS-PAGE and Western blotting analysis.

### Cycloheximide-Chase Assay

HEK293T cells transiently transfected with α1β2γ2(WT) GABA_A_ receptors or stably expressing α1β2γ2(R177G) GABA_A_ receptors were seeded in 6-well plates and incubated at 37 °C overnight. Cells were then treated with 10 μM compound for 24h prior the commencement of the experiment. To stop protein translation, cells were treated with 100 μg/mL cycloheximide (Enzo Life Sciences #ALX-380-269). Cells were then chased for the indicated times, harvested, and lysed for protein analysis.

### Electrophysiological Recordings in Transfected Hek Cells by Whole-Cell Patch Clamp

GABA_A_ α1β2γ2 (WT or R177G) subunits were transfected 48h prior recordings at a subunit plasmid ratio of 0.5α:0.5β:1γ (total DNA was 0.25-0.5 μg) into HEK293T cells using TransIT-2020 reagent (Mirus Bio #MIR 5406, Madison, WI). The α1(WT) subunit was co-transfected with GFP, which acted as an expression marker. ATF6 activators at 10 µM were added to the cultures 24h post-transfection, and 24h prior to the commencement of the experiments.

All experiments were performed at room temperature (RT) in the whole-cell configuration obtained in voltage-clamp mode at a holding potential of -60 mV. Patch pipettes with a resistance of 3 to 5 MΩ were made from borosilicate glass (DWK Life Sciences, #34500-99) on a two-step micropipette puller (Narishige PP-830). The intracellular solution contained the following (in mM): 153 KCl, 1 MgCl_2_, 5 (ethylene glycolbis(b-aminoethyl ether)-N,N,N9,N9-tetraacetic acid (EGTA), 10 4-(2-hydroxyethyl)-1-piperazineethanesulfonic acid (HEPES), and 5 Mg-ATP with pH adjusted to 7.3 and osmolarity at 310 mOsm. Cells were continuously perfused with an extracellular solution composed of (in mM): 142 NaCl, 8 KCl, 6 MgCl_2_, 1 CaCl_2_, 10 Glucose, and 10 HEPES with pH adjusted to 7.4 and osmolarity at 290 mOsm.

Each experiment constituted of one minute recording in extracellular solution with the application of 10 mM GABA (Sigma Aldrich #A2129, Saint Louis, MO) for 4 seconds every 6 seconds to the recorded cell via a theta glass (Siskiyou #15700000E) controlled by a high-speed piezo solution switcher (Siskiyou, MXPZT-300 series).

Evoked inhibitory post-synaptic currents (eIPSCs) were recorded using an Axopatch 200B amplifier and pClamp 10 software, filtered at 1 kHz and sampled at 20 kHz. The first action potential of each recording was analyzed for peak amplitude (pA) using Clampfit 11.2. Normalization of currents to each cell’s capacitance (pF) was performed to allow for collection of current density data (pA/pF). Recordings with an access resistance > 12 MΩ were not included in the analysis. For each experiment, GFP fluorescence was used to control for successful expression of the GABA_A_ receptor in the cells.

### Quantification and Statistical Analysis

Statistical analyses were performed using GraphPad Prism 10 (GraphPad, San Diego, CA). All data were first tested for a Gaussian distribution using a Shapiro-Wilk test. The statistical significance between means was determined by either parametric or nonparametric analysis of variance followed by post-hoc comparisons (Dunn’s, Dunnet) using GraphPad Prism 10. Statistical significance was set at a < 0.05. Error bars in the graphs represent mean ± SEM.

## SUPPORTING INFORMATION INCLUDED

***Supplemental Materials and Methods.*** Document describing the synthesis and characterization of AA263 analogs discussed in this manuscript.

**Table S1**. **RNAseq analysis of HEK293 cells treated with AA263 (10 µM), AA263^yne^ (10 µM), or AA263-5 (10 µM).** There are four sheets in this Excel workbook. DESEQ outputs for RNAseq in HEK293 cells treated with 10 µM AA263, AA263^yne^, or AA263-5 for 6 h and a sheet showing the geneset profiling of different stress-responsive signaling pathways from these RNAseq data.

**Table S2. GO analysis of RNAseq from HEK293 cells treated with AA263 (10 µM), AA263^yne^ (10 µM), or AA263-5 (10 µM).** There are three sheets in this Excel workbook showing the GO analysis results of RNAseq data from HEK293 cells treated for 6 h with 10 µM AA263, AA263^yne^, or AA263-5. GO analysis was performed on genes induced greater than 1.5-fold with an adjusted p value less than 0.05.

## ACKNOWLEDGEMENTS.

We thank Leonard Yoon for help in analyzing the RNAseq experiments described in this manuscript. This work was supported by the National Institutes of Health (AG046495 to RLW and JWK; DK137470 to RLW; NS105789 to TM; HL169631 to WEB).

## COMPETING INTEREST STATEMENT

JWK and RLW are shareholders and scientific advisory board members of Protego Biopharma, which has licensed ATF6 activators for therapeutic development. TM is a scientific advisory board member of Cure GABA- A Variants Foundation, a nonprofit organization.

---

**Figure S1 (Supplement to Figure 1).**
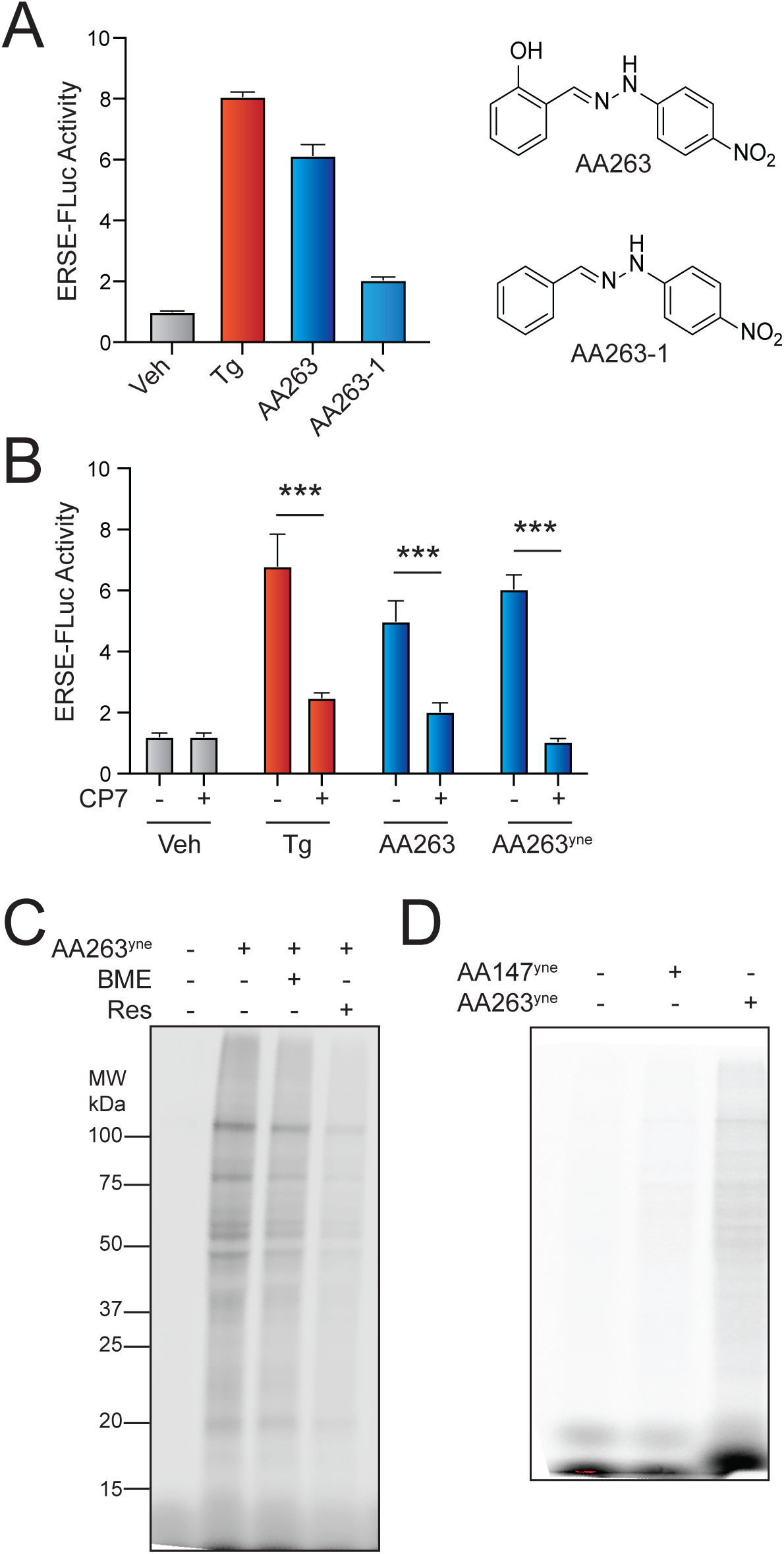
AA263 covalently modifies protein disulfide isomerase family members. **A.** Activation of the ERSE.FLuc ATF6 reporter in HEK293T cells treated for 18 h with vehicle (0.1% DMSO), Tg (500 nM), AA263(10 µM), or AA263-1 (10 μM). Structures of AA263 and AA263-1 shown to the right. Error bars show SEM for n>3 replicates. **B.** Activation of the ERSE.FLuc ATF6 reporter in HEK293T cells treated for 18 h with vehicle (0.1% DMSO), Tg (500 nM), AA263 (10 µM), or AA263^yne^ (10 μM), and/or Ceapin-A7 (CP7; 10 µM). Error bars show SEM for n=3 replicates. ***p<0.005 for two-way ANOVA. **C.** Representative SDS-PAGE gel of Cy5-conjugated proteins from HEK293T cells treated for 4 h with vehicle (0.1% DMSO), AA263^yne^ (10 µM), or the combination of AA263^yne^ (10 µM) and either resveratrol (10 μM) or BME (55 μM). **D.** Representative SDS-PAGE gel of Cy5-conjugated proteins from lysates prepared from HEK293T treated for 1h with vehicle, AA147^yne^, or AA263^yne^ (5 μM).

**Figure S2 (Supplement to Figure 2).**
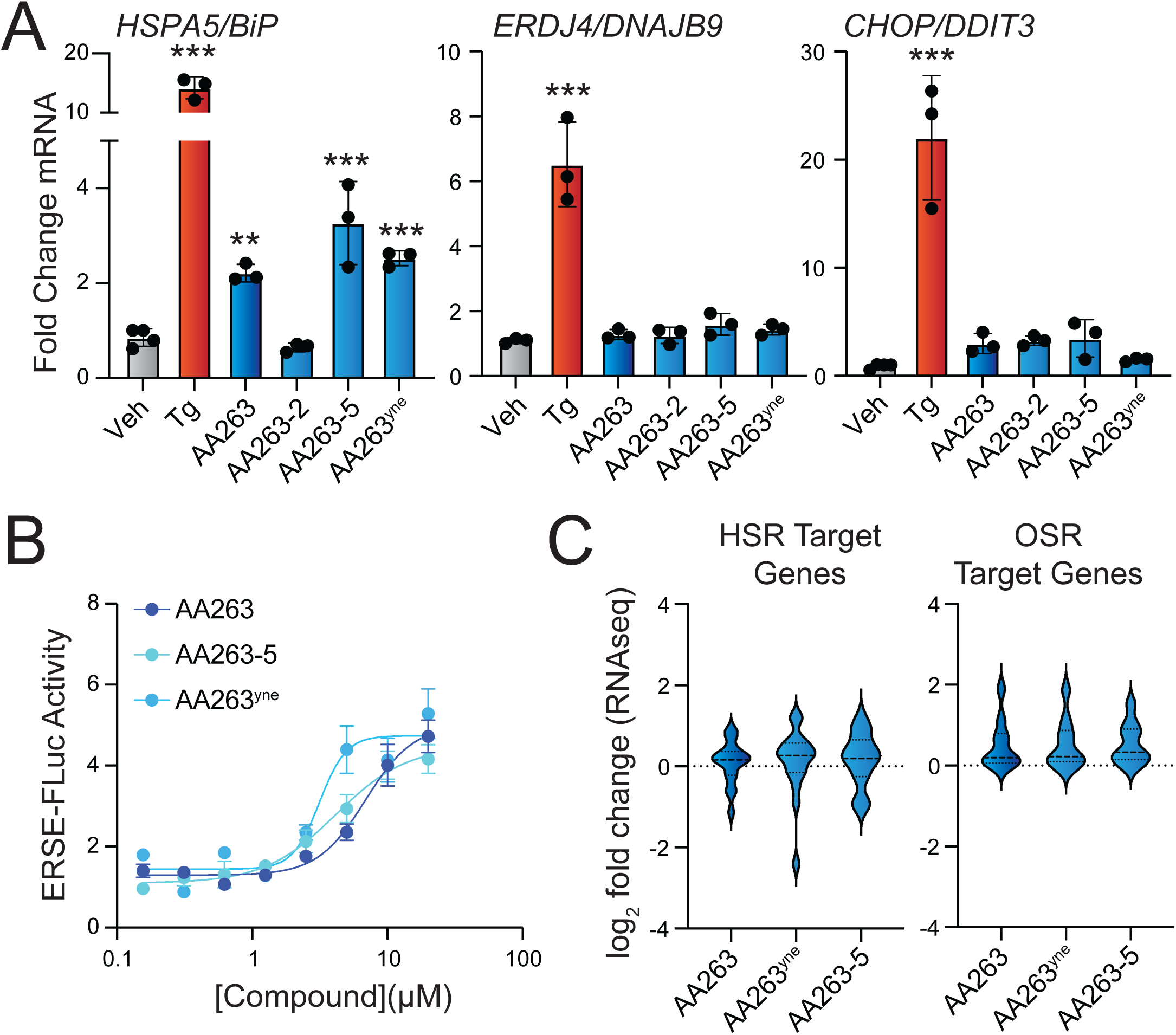
Identification of AA263 analogs that show enhanced ATF6 activation. **A.** Expression, measured by qPCR, of the ATF6 target gene *BiP*, the IRE1/XBP1s target gene *DNAJB9*, and the PERK/ISR target gene *CHOP* in HEK293T cells treated for 6 h with the indicated compound (10 µM). Error bars show SEM for n = 3 biological replicates. *p < 0.05, ***p < 0.001 for one-way ANOVA relative to vehicle-treated cells. **B.** Activation of the ERSE-FLuc ATF6 reporter in HEK293T cells treated for 18 h with the indicated concentration of AA263, AA263^yne^, or AA263-5. Error bars show SEM for n= 3 replicates. The data for AA263 and AA263^yne^ is the same as that shown in Fig. 1E and are included for comparison. **C**. Expression, measured by RNAseq, of gene sets comprising target genes regulated downstream of the heat shock response (HSR, left) or the oxidative stress response (OSR, right) in HEK293 cells treated for 6 h with 10 µM AA263, AA263^yne^, or AA263-5. Full RNAseq data and genesets used in this analysis are shown in **Table S1**

**Figure S3 (Supplement to Figure 3).**
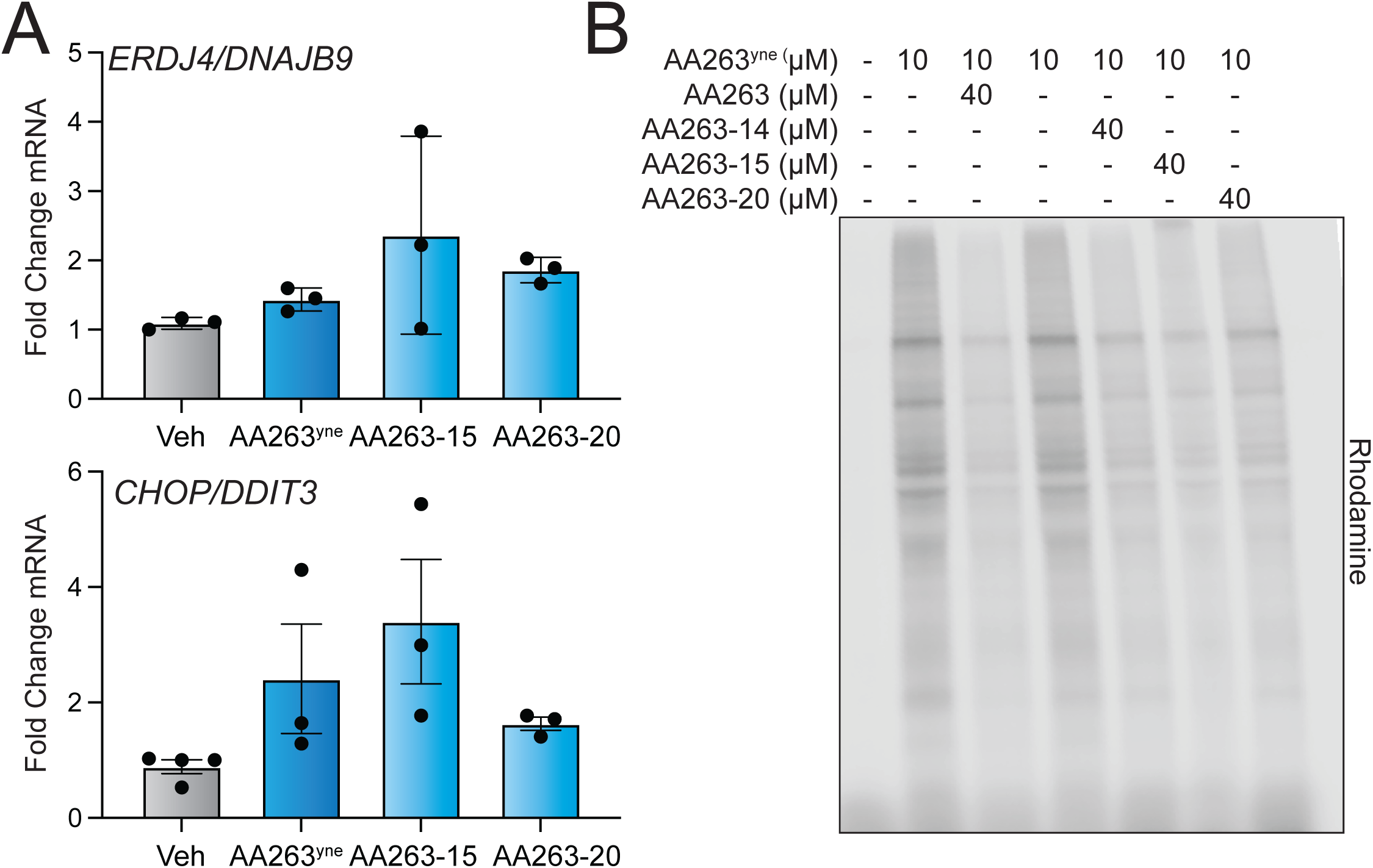
Diversification of the AA263 B-Ring affords improved AA263 analogs. **A.** Expression, measured by qPCR, of the IRE1/XBP1s target gene *DNAJB9* and the PERK/ISR target gene *CHOP* in HEK293T cells treated for 6 h with the indicated AA263 analog (10 µM). Error bars show SEM for n=3 independent biological replicates. **B.** Representative SDS-Page gel of Cy5-conjugated proteins from HEK293T cells treated for 4 h with vehicle, AA263^yne^ (10 μM) or cotreatment of AA263^yne^ (10 μM) with the indicated analog (40 μM).

**Figure S4 (Supplement to Figure 4).**
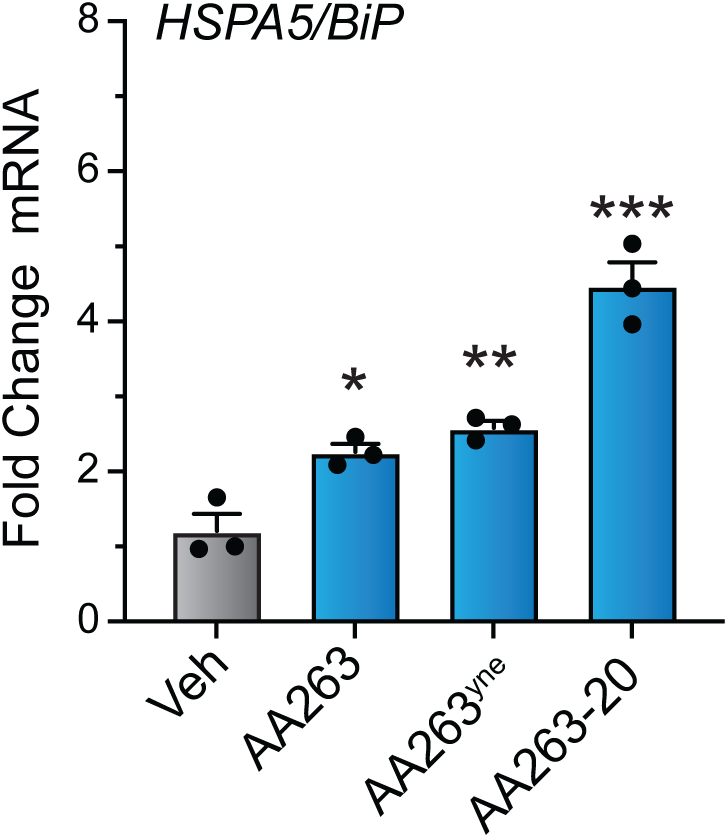
AA263 analogs improve secretory proteostasis for the disease-associated AAT-Z variant. Expression, measured by qPCR, of the ATF6 target gene *BiP* in Huh7.5Z cells treated for 6 h with 10 µM of AA263, AA263^yne^, or AA263-20. Error bars show SEM for n=3 replicates. *p<0.05, **p<0.01, ***p<0.005 for one-way ANOVA compared to vehicle-treated cells are shown.

**Figure S5 (Supplement to Figure 5).**
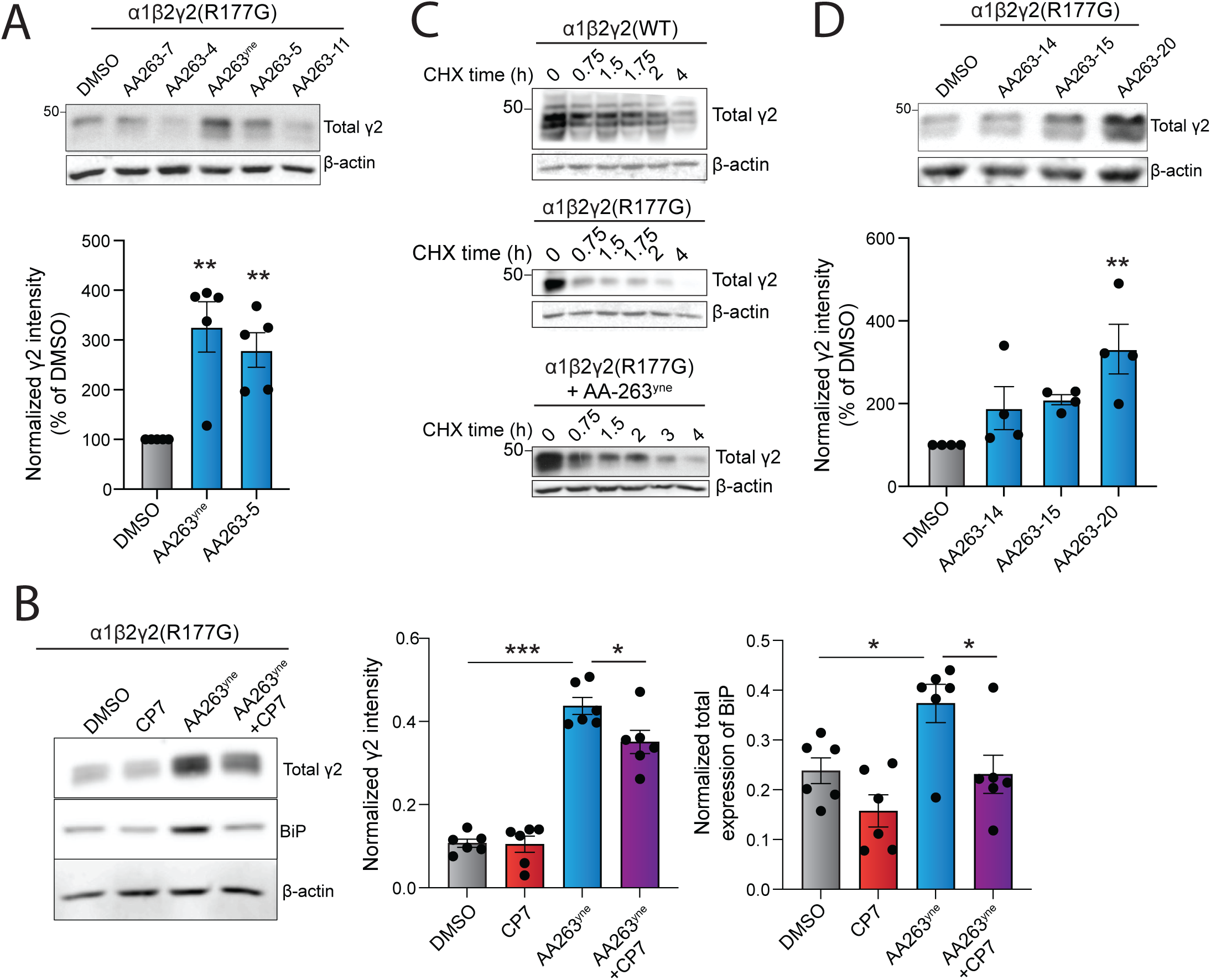
Enhanced AA263 analogs promote the trafficking and plasma membrane activity of destabilized, disease-associated GABA_A_ receptors. **A.** Representative immunoblot (above) of HEK293T cells stably expressing α1β2γ2(R177G) GABA_A_ receptors treated for 24h with the indicated AA263 analog (10 µM). Quantification of band intensities (below) for cells treated with AA263^yne^ or AA263-5 is shown. **B**. Immunoblot and quantification of γ2 and BiP in lysates prepared on HEK293T cells stably expressing α1β2γ2(R177G) GABA_A_ receptors treated for 24h with AA263^yne^ (10 µM) and/or Ceapin-A7 (CP7; 10 µM). **C.** Immunoblot showing the intensity of γ2 over time following cycloheximide (CHX)-chase application (0 to 4 hours, 100 μg/mL) in HEK293T cells transiently transfected with α1β2γ2(WT) receptors (top) and HEK293T cells stably expressing α1β2γ2(R177G) receptor variant treated with vehicle (middle) or AA263^yne^ (bottom). **D**. Representative immunoblot showing total γ2 expression in HEK293T cells stably expressing α1β2γ2(R177G) receptor variant treated with indicated AA263^yne^ analogs (10 µM, 24hrs). One-way ANOVA followed by post-hoc Dunnett’s test was used for statistical analysis. *p≤0.05; **p≤0.01; ****p≤0.0001.

## Notes

### Summary of Updates

This revision addresses comments from the reviewers at ELIFE.

## REFERENCES

1 Hipp, M. S., Kasturi, P. & Hartl, F. U. The proteostasis network and its decline in ageing. Nature Reviews Molecular Cell Biology 20, 421–435, doi:10.1038/s41580-019-0101-y (2019).

2 Jayaraj, G. G., Hipp, M. S. & Hartl, F. U. Functional Modules of the Proteostasis Network. Cold Spring Harbor Perspectives in Biology 12, doi:10.1101/cshperspect.a033951 (2020).

3 Araki, K. & Nagata, K. Protein folding and quality control in the ER. Cold Spring Harbor perspectives in biology 3, a007526 (2011).

4 Kaushik, S. & Cuervo, A. M. Proteostasis and aging. Nature medicine 21, 1406–1415 (2015).

5 Labbadia, J. & Morimoto, R. I. The biology of proteostasis in aging and disease. Annual review of biochemistry 84, 435–464 (2015).

6 Wiseman, R. L., Mesgarzadeh, J. S. & Hendershot, L. M. Reshaping endoplasmic reticulum quality control through the unfolded protein response. Molecular Cell 82, 1477–1491, doi:10.1016/j.molcel.2022.03.025 (2022).

7 Mesgarzadeh, J. S., Buxbaum, J. N. & Wiseman, R. L. Stress-responsive regulation of extracellular proteostasis. Journal of Cell Biology 221, e202112104 (2022).

8 Desport, E., Bridoux, F., Sirac, C., Delbes, S., Bender, S., Fernandez, B., Quellard, N., Lacombe, C., Goujon, J.-M. & Lavergne, D. Al amyloidosis. Orphanet journal of rare diseases 7, 1–13 (2012).

9 Sekijima, Y. Recent progress in the understanding and treatment of transthyretin amyloidosis. Journal of clinical pharmacy and therapeutics 39, 225–233 (2014).

10. Braat, S. & Kooy, R. F. The GABA A Receptor as a Therapeutic Target for Neurodevelopmental Disorders. Neuron **86**, 1119-1130, doi:10.1016/j.neuron.2015.03.042 (2015).

11. Fu, X., Wang, Y. J., Kang, J. Q. & Mu, T. W. in Epilepsy (ed S. J. Czuczwar) (2022).

12 Lomas, D. A. & Mahadeva, R. α 1-Antitrypsin polymerization and the serpinopathies: pathobiology and prospects for therapy. The Journal of clinical investigation 110, 1585–1590 (2002).

13 Silverman, E. K. & Sandhaus, R. A. Alpha1-antitrypsin deficiency. New England Journal of Medicine 360, 2749–2757 (2009).

14 Plate, L. & Wiseman, R. L. Regulating secretory proteostasis through the unfolded protein response: from function to therapy. Trends in cell biology 27, 722–737 (2017).

15 Ron, D. & Walter, P. Signal integration in the endoplasmic reticulum unfolded protein response. Nature reviews Molecular cell biology 8, 519–529 (2007).

16 Walter, P. & Ron, D. The unfolded protein response: from stress pathway to homeostatic regulation. science 334, 1081–1086 (2011).

17 Preissler, S. & Ron, D. Early events in the endoplasmic reticulum unfolded protein response. Cold Spring Harbor perspectives in biology 11, a033894 (2019).

18 Hetz, C., Zhang, K. & Kaufman, R. J. Mechanisms, regulation and functions of the unfolded protein response. Nature reviews Molecular cell biology 21, 421–438 (2020).

19 Adachi, Y., Yamamoto, K., Okada, T., Yoshida, H., Harada, A. & Mori, K. ATF6 is a transcription factor specializing in the regulation of quality control proteins in the endoplasmic reticulum. Cell structure and function 33, 75–89 (2008).

20 Yoshida, H., Matsui, T., Yamamoto, A., Okada, T. & Mori, K. XBP1 mRNA is induced by ATF6 and spliced by IRE1 in response to ER stress to produce a highly active transcription factor. Cell 107, 881–891 (2001).

21 Adamson, B., Norman, T. M., Jost, M., Cho, M. Y., Nuñez, J. K., Chen, Y., Villalta, J. E., Gilbert, L. A., Horlbeck, M. A. & Hein, M. Y. A multiplexed single-cell CRISPR screening platform enables systematic dissection of the unfolded protein response. Cell 167, 1867–1882. e1821 (2016).

22 Shoulders, M. D., Ryno, L. M., Genereux, J. C., Moresco, J. J., Tu, P. G., Wu, C., Yates, J. R., Su, A. I., Kelly, J. W. & Wiseman, R. L. Stress-independent activation of XBP1s and/or ATF6 reveals three functionally diverse ER proteostasis environments. Cell reports 3, 1279–1292 (2013).

23 Chen, J. J., Genereux, J. C., Qu, S., Hulleman, J. D., Shoulders, M. D. & Wiseman, R. L. ATF6 Activation Reduces the Secretion and Extracellular Aggregation of Destabilized Variants of an Amyloidogenic Protein. Chemistry & Biology 21, 1564–1574, doi:10.1016/j.chembiol.2014.09.009 (2014).

24 Cooley, C. B., Ryno, L. M., Plate, L., Morgan, G. J., Hulleman, J. D., Kelly, J. W. & Wiseman, R. L. Unfolded protein response activation reduces secretion and extracellular aggregation of amyloidogenic immunoglobulin light chain. Proceedings of the National Academy of Sciences 111, 13046–13051 (2014).

25 Plate, L., Rius, B., Nguyen, B., Genereux, J. C., Kelly, J. W. & Wiseman, R. L. Quantitative interactome proteomics reveals a molecular basis for ATF6-dependent regulation of a destabilized amyloidogenic protein. Cell chemical biology 26, 913–925. e914 (2019).

26 Plate, L., Cooley, C. B., Chen, J. J., Paxman, R. J., Gallagher, C. M., Madoux, F., Genereux, J. C., Dobbs, W., Garza, D. & Spicer, T. P. Small molecule proteostasis regulators that reprogram the ER to reduce extracellular protein aggregation. elife **5**, e15550 (2016).

27 Paxman, R., Plate, L., Blackwood, E. A., Glembotski, C., Powers, E. T., Wiseman, R. L. & Kelly, J. W. Pharmacologic ATF6 activating compounds are metabolically activated to selectively modify endoplasmic reticulum proteins. Elife 7, e37168 (2018).

28 Higa, A., Taouji, S., Lhomond, S., Jensen, D., Fernandez-Zapico, M. E., Simpson, J. C., Pasquet, J.-M., Schekman, R. & Chevet, E. Endoplasmic reticulum stress-activated transcription factor ATF6α requires the disulfide isomerase PDIA5 to modulate chemoresistance. Molecular and cellular biology 34, 1839–1849 (2014).

29 Koba, H., Jin, S., Imada, N., Ishikawa, T., Ninagawa, S., Okada, T., Sakuma, T., Yamamoto, T. & Mori, K. Reinvestigation of disulfide-bonded oligomeric forms of the unfolded protein response transducer ATF6. Cell Structure and Function 45, 9–21 (2020).

30 Oka, O. B., van Lith, M., Rudolf, J., Tungkum, W., Pringle, M. A. & Bulleid, N. J. ER p18 regulates activation of ATF 6α during unfolded protein response. The EMBO journal 38, e100990 (2019).

31 Oka, O. B. V., Pierre, A. S., Pringle, M. A., Tungkum, W., Cao, Z., Fleming, B. & Bulleid, N. J. Activation of the UPR sensor ATF6α is regulated by its redox-dependent dimerization and ER retention by ERp18. Proceedings of the National Academy of Sciences 119, e2122657119 (2022).

32 Blackwood, E. A., Azizi, K., Thuerauf, D. J., Paxman, R. J., Plate, L., Kelly, J. W., Wiseman, R. L. & Glembotski, C. C. Pharmacologic ATF6 activation confers global protection in widespread disease models by reprograming cellular proteostasis. Nature Communications 10, doi:10.1038/s41467-018-08129-2 (2019).

33 Sun, S., Wang, C., Zhao, P., Kline, G. M., Grandjean, J. M., Jiang, X., Labaudiniere, R., Wiseman, R. L., Kelly, J. W. & Balch, W. E. Capturing the conversion of the pathogenic alpha-1-antitrypsin fold by ATF6 enhanced proteostasis. Cell chemical biology 30, 22–42. e25 (2023).

34 Wang, M., Cotter, E., Wang, Y.-J., Fu, X., Whittsette, A. L., Lynch, J. W., Wiseman, R. L., Kelly, J. W., Keramidas, A. & Mu, T.-W. Pharmacological activation of ATF6 remodels the proteostasis network to rescue pathogenic GABAA receptors. Cell & bioscience 12, 48 (2022).

35 Rius, B., Mesgarzadeh, J. S., Romine, I. C., Paxman, R. J., Kelly, J. W. & Wiseman, R. L. Pharmacologic targeting of plasma cell endoplasmic reticulum proteostasis to reduce amyloidogenic light chain secretion. Blood Advances 5, 1037–1049 (2021).

36 Rosarda, J. D., Stanton, C. R., Chen, E. B., Bollong, M. J. & Wiseman, R. L. Pharmacologic targeting of PDIA1 inhibits NLRP3 Inflammasome assembly and activation. Israel Journal of Chemistry, e202300125 (2023).

37 Kline, G. M., Madrazo, N., Cole, C. M., Pannikkat, M., Bollong, M. J., Rosarda, J. D., Kelly, J. W. & Wiseman, R. L. Metabolically activated proteostasis regulators that protect against erastin-induced ferroptosis. RSC Chemical Biology 5, 866–876 (2024).

38 Kline, G. M., Paxman, R. J., Lin, C.-Y., Madrazo, N., Yoon, L., Grandjean, J. M., Lee, K., Nugroho, K., Powers, E. T. & Wiseman, R. L. Divergent Proteome Reactivity Influences Arm-Selective Activation of the Unfolded Protein Response by Pharmacological Endoplasmic Reticulum Proteostasis Regulators. ACS Chemical Biology 18, 1719–1729 (2023).

39 Ifa, D., Rodrigues, C., De Alencastro, R., Fraga, C. & Barreiro, E. A possible molecular mechanism for the inhibition of cysteine proteases by salicylaldehyde N-acylhydrazones and related compounds. Journal of Molecular Structure: THEOCHEM 505, 11–17 (2000).

40 Torres, S. E., Gallagher, C. M., Plate, L., Gupta, M., Liem, C. R., Guo, X., Tian, R., Stroud, R. M., Kampmann, M., Weissman, J. S. & Walter, P. Ceapins block the unfolded protein response sensor ATF6alpha by inducing a neomorphic inter-organelle tether. Elife 8, doi:10.7554/eLife.46595 (2019).

41 Gallagher, C. M. & Walter, P. Ceapins inhibit ATF6alpha signaling by selectively preventing transport of ATF6alpha to the Golgi apparatus during ER stress. Elife 5, doi:10.7554/eLife.11880 (2016).

42 Gallagher, C. M., Garri, C., Cain, E. L., Ang, K. K., Wilson, C. G., Chen, S., Hearn, B. R., Jaishankar, P., Aranda-Diaz, A., Arkin, M. R., Renslo, A. R. & Walter, P. Ceapins are a new class of unfolded protein response inhibitors, selectively targeting the ATF6alpha branch. Elife 5, doi:10.7554/eLife.11878 (2016).

43 Chan, W. C., Sharifzadeh, S., Buhrlage, S. J. & Marto, J. A. Chemoproteomic methods for covalent drug discovery. Chemical Society Reviews 50, 8361–8381 (2021).

44 Zhang, L. & Elias, J. E. Relative protein quantification using tandem mass tag mass spectrometry. Proteomics: methods and protocols, 185–198 (2017).

45 Oka, O. B. V., Pierre, A. S., Pringle, M. A., Tungkum, W., Cao, Z., Fleming, B. & Bulleid, N. J. Activation of the UPR sensor ATF6alpha is regulated by its redox-dependent dimerization and ER retention by ERp18. Proc Natl Acad Sci U S A 119, e2122657119, doi:10.1073/pnas.2122657119 (2022).

46 Oka, O. B., van Lith, M., Rudolf, J., Tungkum, W., Pringle, M. A. & Bulleid, N. J. ERp18 regulates activation of ATF6alpha during unfolded protein response. EMBO J 38, e100990, doi:10.15252/embj.2018100990 (2019).

47 Backus, K. M., Correia, B. E., Lum, K. M., Forli, S., Horning, B. D., Gonzalez-Paez, G. E., Chatterjee, S., Lanning, B. R., Teijaro, J. R., Olson, A. J., Wolan, D. W. & Cravatt, B. F. Proteome-wide covalent ligand discovery in native biological systems. Nature 534, 570–574, doi:10.1038/nature18002 (2016).

48 Paxman, R., Plate, L., Blackwood, E. A., Glembotski, C., Powers, E. T., Wiseman, R. L. & Kelly, J. W. Pharmacologic ATF6 activating compounds are metabolically activated to selectively modify endoplasmic reticulum proteins. Elife 7, doi:10.7554/eLife.37168 (2018).

49 Kline, G. M., Paxman, R. J., Lin, C. Y., Madrazo, N., Yoon, L., Grandjean, J. M. D., Lee, K., Nugroho, K., Powers, E. T., Wiseman, R. L. & Kelly, J. W. Divergent Proteome Reactivity Influences Arm-Selective Activation of the Unfolded Protein Response by Pharmacological Endoplasmic Reticulum Proteostasis Regulators. ACS Chem Biol 18, 1719–1729, doi:10.1021/acschembio.3c00042 (2023).

50 Coe, K. J., Jia, Y., Ho, H. K., Rademacher, P., Bammler, T. K., Beyer, R. P., Farin, F. M., Woodke, L., Plymate, S. R. & Fausto, N. Comparison of the cytotoxicity of the nitroaromatic drug flutamide to its cyano analogue in the hepatocyte cell line TAMH: evidence for complex I inhibition and mitochondrial dysfunction using toxicogenomic screening. Chemical research in toxicology 20, 1277–1290 (2007).

51 Nepali, K., Lee, H.-Y. & Liou, J.-P. Nitro-group-containing drugs. Journal of medicinal chemistry 62, 2851–2893 (2018).

52 Grandjean, J. M., Plate, L., Morimoto, R. I., Bollong, M. J., Powers, E. T. & Wiseman, R. L. Deconvoluting stress-responsive proteostasis signaling pathways for pharmacologic activation using targeted RNA sequencing. ACS chemical biology 14, 784–795 (2019).

53 Lebeau, J., Saunders, J. M., Moraes, V. W., Madhavan, A., Madrazo, N., Anthony, M. C. & Wiseman, R. L. The PERK arm of the unfolded protein response regulates mitochondrial morphology during acute endoplasmic reticulum stress. Cell reports 22, 2827–2836 (2018).

54 Verfaillie, T., Rubio, N., Garg, A., Bultynck, G., Rizzuto, R., Decuypere, J., Piette, J., Linehan, C., Gupta, S. & Samali, A. PERK is required at the ER-mitochondrial contact sites to convey apoptosis after ROS-based ER stress. Cell Death & Differentiation 19, 1880–1891 (2012).

55 Mi, H., Muruganujan, A., Casagrande, J. T. & Thomas, P. D. Large-scale gene function analysis with the PANTHER classification system. Nature protocols 8, 1551–1566 (2013).

56 Bieth, J., Spiess, B. & Wermuth, C. G. The synthesis and analytical use of a highly sensitive and convenient substrate of elastase. Biochemical medicine 11, 350–357 (1974).

57 Braat, S. & Kooy, R. F. The GABAA receptor as a therapeutic target for neurodevelopmental disorders. Neuron 86, 1119–1130 (2015).

58 Sigel, E. & Steinmann, M. E. Structure, function, and modulation of GABAA receptors. Journal of Biological Chemistry 287, 40224–40231 (2012).

59 Benarroch, E. E. GABAA receptor heterogeneity, function, and implications for epilepsy. Neurology 68, 612–614 (2007).

60 Wang, Y. J., Seibert, H., Ahn, L. Y., Schaffer, A. E. & Mu, T. W. Pharmacological chaperones restore proteostasis of epilepsy-associated GABA(A) receptor variants. Pharmacol Res 208, 107356, doi:10.1016/j.phrs.2024.107356 (2024).

61 Todd, E., Gurba, K. N., Botzolakis, E. J., Stanic, A. K. & Macdonald, R. L. GABAA receptor biogenesis is impaired by the γ2 subunit febrile seizure-associated mutation, GABRG2 (R177G). Neurobiology of disease 69, 215–224 (2014).

62 Di, X.-J., Wang, Y.-J., Cotter, E., Wang, M., Whittsette, A. L., Han, D.-Y., Sangwung, P., Brown, R., Lynch, J. W. & Keramidas, A. Proteostasis regulators restore function of epilepsy-associated GABAA receptors. Cell chemical biology 28, 46–59. e47 (2021).

63 Todd, E., Gurba, K. N., Botzolakis, E. J., Stanic, A. K. & Macdonald, R. L. GABAA receptor biogenesis is impaired by the gamma2 subunit febrile seizure-associated mutation, GABRG2(R177G). Neurobiol Dis 69, 215-224, doi:10.1016/j.nbd.2014.05.013 (2014).

64 Wang, M., Cotter, E., Wang, Y. J., Fu, X., Whittsette, A. L., Lynch, J. W., Wiseman, R. L., Kelly, J. W., Keramidas, A. & Mu, T. W. Pharmacological activation of ATF6 remodels the proteostasis network to rescue pathogenic GABA(A) receptors. Cell Biosci 12, 48, doi:10.1186/s13578-022-00783-w (2022).

65 Ifa, D. R., Rodrigues, C. R., de Alencastro, R. B., Fraga, C. A. M. & Barreiro, E. J. A possible molecular mechanism for the inhibition of cysteine proteases by salicylaldehyde *N*-acylhydrazones and related compounds. Journal of Molecular Structure-Theochem 505, 11–17, doi:10.1016/s0166-1280(99)00307-3 (2000).

66 Zambaldo, C., Vinogradova, E. V., Qi, X. T., Iaconelli, J., Suciu, R. M., Koh, M., Senkane, K., Chadwick, S. R., Sanchez, B. B., Chen, J. S., Chatterjee, A. K., Liu, P., Schultz, P. G., Cravatt, B. F. & Bollong, M. J. 2-Sulfonylpyridines as Tunable, Cysteine-Reactive Electrophiles. Journal of the American Chemical Society 142, 8972–8979, doi:10.1021/jacs.0c02721 (2020).

67 Shindo, N., Fuchida, H., Sato, M., Watari, K., Shibata, T., Kuwata, K., Miura, C., Okamoto, K., Hatsuyama, Y., Tokunaga, K., Sakamoto, S., Morimoto, S., Abe, Y., Shiroishi, M., Caaveiro, J. M. M., Ueda, T., Tamura, T., Matsunaga, N., Nakao, T., Koyanagi, S., Ohdo, S., Yamaguchi, Y., Hamachi, I., Ono, M. & Ojida, A. Selective and reversible modification of kinase cysteines with chlorofluoroacetamides. Nature Chemical Biology 15, 250-+, doi:10.1038/s41589-018-0204-3 (2019).

68 Higa, A., Taouji, S., Lhomond, S., Jensen, D., Fernandez-Zapico, M. E., Simpson, J. C., Pasquet, J. M., Schekman, R. & Chevet, E. Endoplasmic reticulum stress-activated transcription factor ATF6alpha requires the disulfide isomerase PDIA5 to modulate chemoresistance. Mol Cell Biol 34, 1839–1849, doi:10.1128/MCB.01484-13 (2014).

69 Eletto, D., Eletto, D., Dersh, D., Gidalevitz, T. & Argon, Y. Protein Disulfide Isomerase A6 Controls the Decay of IRE1α Signaling via Disulfide-Dependent Association. Molecular Cell 53, 562–576, doi:10.1016/j.molcel.2014.01.004 (2014).

70 Kranz, P., Neumann, F., Wolf, A., Classen, F., Pompsch, M., Ocklenburg, T., Baumann, J., Janke, K., Baumann, M., Goepelt, K., Riffkin, H., Metzen, E. & Brockmeier, U. PDI is an essential redox-sensitive activator of PERK during the unfolded protein response (UPR). Cell Death & Disease 8, doi:10.1038/cddis.2017.369 (2017).

71 Coe, K. J., Jia, Y., Ho, H. K., Rademacher, P., Bammler, T. K., Beyer, R. P., Farin, F. M., Woodke, L., Plymate, S. R., Fausto, N. & Nelson, S. D. Comparison of the cytotoxicity of the nitroaromatic drug flutamide to its cyano analogue in the hepatocyte cell line TAMH: Evidence for complex I inhibition and mitochondrial dysfunction using toxicogenomic screening. Chemical Research in Toxicology 20, 1277–1290, doi:10.1021/tx7001349 (2007).

72 Nepali, K., Lee, H. Y. & Liou, J. P. Nitro-Group-Containing Drugs. Journal of Medicinal Chemistry 62, 2851–2893, doi:10.1021/acs.jmedchem.8b00147 (2019).

73 Bassot, A., Chen, J. S., Takahashi-Yamashiro, K., Yap, M. C., Gibhardt, C. S., Le, G. N. T., Hario, S., Nasu, Y., Moore, J., Gutierrez, T., Mina, L., Mast, H., Moses, A., Bhat, R., Ballanyi, K., Lemieux, H., Sitia, R., Zito, E., Bogeski, I., Campbell, R. E. & Simmen, T. The endoplasmic reticulum kinase PERK interacts with the oxidoreductase ERO1 to metabolically adapt mitochondria. Cell Reports 42, doi:10.1016/j.celrep.2022.111899 (2023).

74 Lebeau, J., Saunders, J. M., Moraes, V. W. R., Madhavan, A., Madrazo, N., Anthony, M. C. & Wiseman, R. L. The PERK Arm of the Unfolded Protein Response Regulates Mitochondrial Morphology during Acute Endoplasmic Reticulum Stress. Cell Reports 22, 2827–2836, doi:10.1016/j.celrep.2018.02.055 (2018).

75 Verfaillie, T., Rubio, N., Garg, A. D., Bultynck, G., Rizzuto, R., Decuypere, J. P., Piette, J., Linehan, C., Gupta, S., Samali, A. & Agostinis, P. PERK is required at the ER-mitochondrial contact sites to convey apoptosis after ROS-based ER stress. Cell Death and Differentiation 19, 1880–1891, doi:10.1038/cdd.2012.74 (2012).

76 Baell, J. B. & Holloway, G. A. New Substructure Filters for Removal of Pan Assay Interference Compounds (PAINS) from Screening Libraries and for Their Exclusion in Bioassays. Journal of Medicinal Chemistry 53, 2719–2740, doi:10.1021/jm901137j (2010).

77 Baell, J. B. & Nissink, J. W. M. Seven Year Itch: Pan-Assay Interference Compounds (PAINS) in 2017- Utility and Limitations. Acs Chemical Biology 13, 36–44, doi:10.1021/acschembio.7b00903 (2018).

78 Pouliot, M. & Jeanmart, S. Pan Assay Interference Compounds (PAINS) and Other Promiscuous Compounds in Antifungal Research. Journal of Medicinal Chemistry 59, 497–503, doi:10.1021/acs.jmedchem.5b00361 (2016).

79 Park, B. K., Boobis, A., Clarke, S., Goldring, C. E. P., Jones, D., Kenna, J. G., Lambert, C., Laverty, H. G., Naisbitt, D. J., Nelson, S., Nicoll-Griffith, D. A., Obach, R. S., Routledge, P., Smith, D. A., Tweedie, D. J., Vermeulen, N., Williams, D. P., Wilson, I. D. & Baillie, T. A. Managing the challenge of chemically reactive metabolites in drug development. Nature Reviews Drug Discovery 10, 292–306, doi:10.1038/nrd3408 (2011).

80 Plate, L., Cooley, C. B., Chen, J. J., Paxman, R. J., Gallagher, C. M., Madoux, F., Genereux, J. C., Dobbs, W., Garza, D., Spicer, T. P., Scampavia, L., Brown, S. J., Rosen, H., Powers, E. T., Walter, P., Hodder, P., Wiseman, R. L. & Kelly, J. W. Small molecule proteostasis regulators that reprogram the ER to reduce extracellular protein aggregation. Elife 5, 1–26, doi:10.7554/eLife.15550 (2016).

81 Grandjean, J. M. D. & Wiseman, R. L. Small molecule strategies to harness the unfolded protein response: where do we go from here? J Biol Chem 295, 15692–15711, doi:10.1074/jbc.REV120.010218 (2020).

82 Lu, Y., Wang, L. R., Lee, J., Mohammad, N. S., Aranyos, A. M., Gould, C., Khodayari, N., Oshins, R. A., Moneypenny, C. G. & Brantly, M. L. The unfolded protein response to PI*Z alpha-1 antitrypsin in human hepatocellular and murine models. Hepatol Commun 6, 2354–2367, doi:10.1002/hep4.1997 (2022).

83 Wang, C., Zhao, P., Sun, S., Wang, X. & Balch, W. (bioRxiv, 2022).

84 Miranda, E., Pérez, J., Ekeowa, U. I., Hadzic, N., Kalsheker, N., Gooptu, B., Portmann, B., Belorgey, D., Hill, M., Chambers, S., Teckman, J., Alexander, G. J., Marciniak, S. J. & Lomas, D. A. A novel monoclonal antibody to characterize pathogenic polymers in liver disease associated with alpha1- antitrypsin deficiency. Hepatology 52, 1078–1088, doi:10.1002/hep.23760 (2010).

